# Experimental Disturbance-Induced Shifts in Benthic Functional Diversity: the importance of marine protected areas in soft-bottom ecosystems

**DOI:** 10.64898/2026.06.01.729212

**Authors:** Kasper J. Meijer, Oscar Franken, Laura L. Govers, Tjisse van der Heide, Eva Willebrands, Mark E. de Wilt, Bas de Wit, Han Olff

## Abstract

Benthic communities form the foundation of food webs in coastal and shelf seas worldwide. However, these communities face severe threats from anthropogenic bottom-disturbing activities. Marine Protected Areas (MPAs) effectively safeguard benthic communities against such disturbances. Nevertheless, the effectiveness of MPAs for the recovery of benthic communities in highly dynamic soft-bottom areas is much debated, as these communities are thought to be adapted to natural dynamics and therefore resistant to human impacts. In this study, we examine whether MPAs in highly dynamic seas support sensitive communities that are affected by bottom-disturbing fishing gear. Utilising a Before-After-Control-Impact (BACI) experimental design, we investigated the effects of disturbance caused by a shrimp trawl gear on macrozoobenthic communities within MPAs in a dynamic coastal soft-bottom sea over two years. We assessed changes in taxonomic and functional diversity, as well as community composition, across three levels of trawling intensity and frequency (low intensity and high frequency, high intensity and low frequency, and high intensity and high frequency) compared with undisturbed control areas. The study was carried out in the lower dynamic areas of the study system, where we found that, even light trawling gear impacted functional diversity and community assembly of macrozoobenthic communities, leading to shifts in functional groups. Contrastingly, taxonomic diversity metrics were not affected. We found that taxonomic and functional turnover increased stronger over time than the undisturbed control, with the highest turnover in the most frequently disturbed treatments. Long-living species were negatively impacted by disturbance, and the abundance of scavengers and predators increased in trawled areas, with possible radiating effects into undisturbed areas, while suspension feeders declined. Our findings demonstrate that anthropogenic disturbance remains a relevant ecological pressure, even in areas that may appear resilient due to their dynamic nature. MPAs can then provide ecological benefits by protecting sensitive and functionally important species.

**Article Impact Statement:** Marine Protected Areas are effective in safeguarding sensitive communities from anthropogenic disturbance.

## Introduction

Human activities have increasingly impacted ecosystems worldwide, with coastal areas experiencing some of the highest anthropogenic pressures (Harley et al., 2006; Halpern et al., 2008; 2019; He & Silliman, 2019). Coastal systems are among the most productive systems globally, with critical ecosystem services such as carbon sequestration, nutrient cycling, and maintaining fish stocks (Barbier et al., 2011). However, this also makes them highly attractive for fisheries and other activities, exacerbating conflicts between conservation and socioeconomic interests (Halpern et al., 2008; Barbier et al., 2011). Marine Protected Areas are among the most effective conservation strategies for minimising human impacts and protecting biodiversity (Dudley, 2008; Rising & Heal, 2014; Sala et al., 2018). However, the effectiveness of Marine Protected Areas is often compromised by human activities that are sometimes still allowed within their boundaries (Lubchenco & Grorud-Colvert, 2015; Sala & Giakoumi, 2018; Zupan et al., 2018). Human activities with little to no impact can be permitted within designated Marine Protected Areas without affecting the ecological integrity of the system (Grorud-Colvert et al., 2021). Assessing the effectiveness of Marine Protected Areas in preserving ecological integrity by prohibiting detrimental human activities requires a comprehensive understanding of how these activities alter ecological functioning and the contexts in which this occurs.

Context-dependent relationships between ecological functioning and habitat type are important considerations for management (Bradly et al. 2020). Quantifying the extent to which anthropogenic activities disturb seafloor integrity is important for generalising findings across systems and for applying effective conservation measures. Benthic communities in dynamic soft-bottom systems are naturally adapted to frequent physical disturbances, such as strong currents and sediment movement (Kaiser et al., 2002). These communities typically consist of species with early maturation, fast growth rates, and small sizes, whereas in more calm environments, larger, long-lived and slow-growing species dominate (Pianka 1970). Physical disturbance of the seafloor can then shift community composition towards smaller and short-lived species despite low levels of natural dynamics (Hiddink et al., 2006; van Denderen et al., 2015; McLaverty et al., 2024). Such shifts in community structure can have cascading effects on ecosystem functioning, including reductions in bioturbation, nutrient cycling, and organic matter decomposition, processes that are often driven by the activity of larger, longer-lived benthic organisms (Queirós et al., 2013; Solan et al., 2004; Mermillod-Blondin & Rosenberg, 2006). Consequently, the benthic communities found in low-dynamic areas are thought to be particularly vulnerable to physical disturbances (Kindsvater et al., 2016). Conversely, the inherent resilience of communities in high-dynamic environments might suggest that restricting bottom-disturbing activities within Marine Protected Areas in such areas yields limited additional benefit, as communities can recover relatively quickly from low levels of impact (Hiddink et al., 2017; Sciberras et al., 2018). Especially in dynamic coastal systems, such as soft-bottom estuaries, identifying and protecting refuge areas, i.e., areas with relatively low natural dynamic energy, is critical for conserving vulnerable natural values and ecosystem functions. Yet, it remains uncertain how effectively Marine Protected Areas can protect vulnerable habitats and provide ecological benefits in environments where natural forces already play a dominant role in structuring communities.

Evaluating the ecological effectiveness of Marine Protected Areas requires long-term monitoring, with measurable effects sometimes only emerging after years or even decades (Micheli et al., 2004; Grorud-Colvert et al., 2021). However, Marine Protected Areas are often established opportunistically (Rabaut et al., 2009), without the opportunity to implement rigorous baseline monitoring. This lack of pre-closure data limits the ability to conduct statistically robust before–and–after comparisons, complicating evaluations of ecological change. To address this limitation, we employed a reverse experimental approach, reintroducing physical bottom disturbance to previously protected areas to measure ecosystem responses to renewed anthropogenic impact, rather than waiting for long-term recovery. This allows us to assess the effectiveness of protection by observing the impact of reintroducing human activities that had been previously excluded from human activities for six years.

As a model system, we use the Dutch Wadden Sea, a soft-bottom estuarine system where several bottom-disturbing activities are permitted, despite its UNESCO World Heritage Status and Natura 2000 designation (Bastmeijer et al., 2023; Meijer et al., 2025). European Union directives (e.g., the Species and Habitats Directive 92/43/EEC and the Birds Directive 79/409/EEC) allow anthropogenic activities within Natura 2000 areas, provided they do not significantly compromise the ecological objectives of the site. The Natura 2000 management plan outlines the use of area-based protection and restrictions on human activities as key tools for achieving these objectives (Ministerie van Infrastructuur en Milieu, Rijkswaterstaat Noord-Nederland, 2016). As a result, approximately 10% of the subtidal area in the Dutch Wadden Sea is currently fully closed to bottom-disturbing activities (Meijer et al., 2025). However, the Wadden Sea overall is considered a highly dynamic system and thus thought to be more resilient to additional anthropogenic disturbances (Bergman et al., 2015; Rijnsdorp et al., 2018; Holzhauer et al., 2022). Still, varying degrees of dynamics have been classified, with 66% of the sublittoral area classified as low-dynamic (Baptist et al., 2019, 2021), and receiving the majority of efforts in terms of protection (Meijer et al., 2025). Nonetheless, the ecological effectiveness of these Marine Protected Areas in enhancing benthic community integrity remains empirically untested.

In this study, we investigate the response of macrozoobenthic communities to experimentally reintroduced physical disturbance in areas currently protected from bottom-disturbing activities. Specifically, we simulate multiple disturbances by trawling within Marine Protected Areas using shrimp gear without nets. This is to replicate the mechanical impact of fishing gear, with which parallels can be drawn to other disturbance activities, while avoiding confounding effects through the direct removal of organisms. Still, a shrimp gear with a full net will likely have a higher mechanical impact than simulated here due to the net sweeping the seafloor. Using a Before-After-Control-Impact (BACI) experimental design, we independently manipulated the intensity and frequency levels of bottom disturbance in 48-ha plots in Marine Protected Areas to examine how these factors influence benthic community composition and ecosystem functions over time. We do this by investigating the effects of bottom disturbance on 1) taxonomic and functional composition, 2) taxonomic and functional diversity, and 3) dominating functional groups of the benthic communities in the Dutch Wadden Sea. In addition to traditional community analysis, we apply a functional trait approach to identify life history strategies responsive to disturbance, thereby enabling generalisations of our findings across geographic areas where species might be taxonomically distinct but functionally similar. Specifically, we expect to find a decrease in long-living species in impacted areas as these are generally more vulnerable to bottom disturbance (Rijnsdorp et al., 2018), whereas we expect scavenging species to increase due to the enhanced food availability following the direct mortality of organisms caused by passing trawls (Tillin et al., 2006; McLaverty et al., 2024). This study offers crucial insights into whether and how area-based protection mitigates the impacts of human-induced bottom disturbance, even in a naturally dynamic environment, informing future ecosystem-based marine spatial planning and conservation strategies in economically important coastal regions.

## Materials and Methods

### Study site

The study was conducted in the westernmost tidal basin of the Wadden Sea (Figure 1A). This Marsdiep tidal basin (Figure 1B) is the largest tidal basin (690 km^2^) of the Dutch Wadden Sea (Oost et al. 2018). It contains the largest area of subtidal (permanently submerged) habitat (Baptist et al., 2019). Ecotopes have been classified for the Dutch Wadden Sea for management purposes (van Donk & Baptist, 2021). Currently, the subtidal parts of the Marsdiep tidal basin are divided into six ecotopes (Paree et al., 2023) based on depth and dynamics, with the largest area covered by the shallow low dynamic ecotope (Figure 1B). Marine Protected Areas are largely focused in the low-dynamic shallow subtidal which host the highest ecological values and most sensitive communities to bottom disturbing activities (Meijer et al., 2025). Except for anchoring, all bottom-disturbing activities have been prohibited in these areas for six years prior to the start of our study in 2021. We use shrimp fishing to simulate bottom disturbance as this is the most pervasive human activity in the Dutch Wadden Sea (Temming & Hufnagl, 2015; Tulp et al., 2016). This type of fishery uses lighter gear, is towed at lower speeds, and lacks tickler chains, relying instead on a single chain with rollers (Bobbin rope), which results in lower seabed penetration and benthic impact (Hiddink et al., 2017; Tulp et al., 2020). Still, trawling frequencies exceeding 30 times a year are common in the Wadden Sea, possibly impacting benthic communities through repeated and persistent disturbance (ICES, 2018; Tulp et al., 2020).

**Figure 1.**
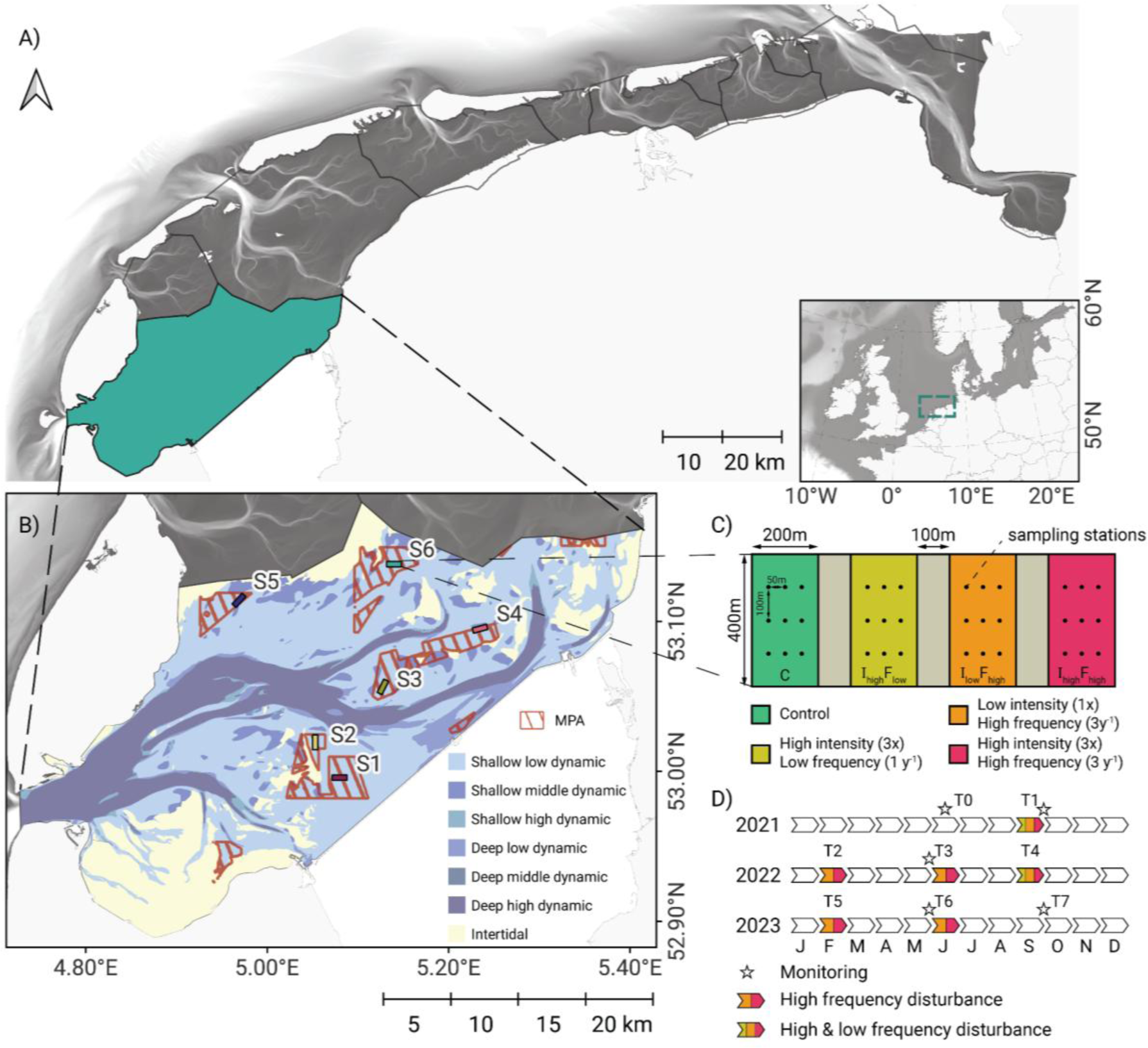
Study site and experimental design. A) Overview of the Dutch Wadden Sea with the tidal basin Marsdiep indicated in green. B) The Marsdiep tidal basin with classified ecotopes and the location of the six study sites. All sites are placed within Marine Protected Areas that have excluded all fisheries since 2015. C) Close-up of each study site and the treatments. At each site, four different disturbance treatments were applied (combinations of disturbance frequency and disturbance intensity). Treatment plots within a site each measured 400×200m and had a 100m buffer zone between treatments. The spatial arrangement of treatments was randomised between sites (see Supplementary Figure S1). Nine fixed sampling locations were used within each treatment. D) Timeline of the disturbances and the monitoring periods from June 2021 until October 2023. The colours indicate which treatments were included in each disturbance.

### Experimental design

Experimental sites are located in different Marine Protected Areas in the Marsdiep tidal basin in the Dutch Wadden Sea (Figure 1B). Each site contains four treatments, each 400×200m, with a buffer zone of 100m between treatments (Figure 1C). Experimental disturbance events were carried out by a commercial shrimp fishing vessel (length 22m, width 6m). The disturbance treatments involved trawling the area following standard shrimp fishing practices at a speed of 2.5 knots, using a shrimp trawl that measured 7 meters on either side of the vessel. No fishing net was attached to the trawl during the experiment as we wanted to specifically test the physical disturbance effect of bottom disturbance, not the trophic impact of catch or bycatch from fishing (which requires evaluation at much larger scales). In doing so, we only look at the main impact of the trawl through its weighted bottom-side chafer that drags across the seafloor, the shoe, and the single chain fitted with rolling bobbin wheels to guide the net over the top layer of the seabed. The treatments consist of a control (C) treatment (no disturbance), high intensity and low frequency (I_high_F_low_) treatment (disturbed once per year with three repeated area-covering trawls per disturbance event, i.e. Swept Area Ratio = 3), low intensity and high frequency (I_low_F_high_) treatment (disturbed three times per year with a single area-covering trawl per disturbance event, i.e. Swept Area Ratio = 3), and a high intensity and high frequency (I_high_F_high_) treatment (disturbed three times per year with three area covering trawls per disturbance event, i.e. Swept Area Ratio = 9) (Figure 1C). We specifically varied intensity and frequency to test whether longer recovery times between disturbance events make up for the intensity of the disturbance (resistance and recovery potential). The order of treatments was randomised between sites (Supplementary Figure S1). Nine fixed sampling locations were located within each treatment, separated by at least 50 m (Figure 1C).

Sampling points were sampled for benthic macrofauna by the NIOZ Research Vessel Navicula, using a 20 x 30 cm (0.06 m2) hydraulic boxcore to a depth of ∼25 cm. The sampling stations were approached by the research vessel with an accuracy of approximately 15m. In combination with the size of the boxcore, it was therefore highly unlikely that exactly the same part of the seafloor was sampled during subsequent sampling events, even though the same location was used. A total of 30 out of 1080 samples were excluded from the analysis as the sampling failed up to three times, resulting in empty or half-filled boxcores (Supplementary Table S1). Each sample was sieved over a 1mm round mesh. Larger shellfish and crustaceans were frozen at -20℃. The remainder of the sample was stored in a borax-buffered 6% formalin:seawater solution with ∼2 mg L^-1^ Rose Bengal (CAS no. 632-68-8). All organisms were sorted, counted, and identified in the lab to the lowest possible taxonomic resolution. Sometimes it was impossible to identify individuals to the species level, especially for juveniles or when individuals were damaged. In case both genus-level identification and species-level identification within the same genus were found in a sample, the abundance of the genus was added to the species level. This was done in the ratio of each species’ occurrence in the sample, when multiple species of the same genus were found. The genus level was used if no individuals from that genus were identified to the species level in the sample.

Experimental plots were sampled 5 times: at the start of the experiment, before the disturbance (T0), directly after the first disturbance event (T1), before the third (T3) and sixth (T6) disturbance event as well as one final time after the sixth disturbance event to measure the recovery (T7), resulting in a Before-After-Control-Impact (BACI) design study (Osenberg et al., 2006; Stewart-Oaten et al., 1986). Sampling before applying any of the treatments (T0) occurred in June 2021, and the first disturbance event occurred in September 2021, in which all treatments except for the control treatment were trawled (Figure 1D). A sampling round immediately followed after this disturbance, to look at the short-term effects of disturbance (i.e., one week) (Figure 1D). Subsequently, sampling occurred one week before the disturbance event in June 2022 and 2023 (Figure 1D). A final sampling round took place in September 2023, two years after the initial disturbance. During the data analysis, the BACI design could account for existing initial differences between the study locations when comparing the impacts of the different disturbance treatments.

### Functional traits

Functional trait data on 15 life history traits were extracted from an existing functional trait database (Gusmao et al., 2022; Meijer et al., 2023). These included adult body size, adult burrowing depth, adult locomotion, age of sexual maturation, bioturbation mode, fecundity, feeding type, larval development location, living habitat, longevity, offspring size, offspring type, reproductive frequency, reproductive mode, and skeleton (Supplementary Table S2). The database contains information on taxonomic groups for several life-history traits, each subdivided into modalities (categories) and scored for affinity using a fuzzy-coding approach (Chevenet et al., 1994), with scores ranging from 0 to 3. A score of zero is given when no affinity is found for a certain modality, whereas three is given when no affinity for any other modality is found. Taxa expressing affinity for multiple modalities are scored with a one or a two, where a one represents lesser affinity, and a two represents greater or equal affinity. Fuzzy scores are standardised to a value between zero and one and multiplied by the abundance of each species in each sample. The community-weighted mean is then determined for each trait modality by dividing by the total number of individuals in each sample.

### Statistical analysis

We used the benthic community data from the boxcore samples in several analyses. First, to get an indication of responses to disturbance-sensitive versus disturbance-tolerant species, we calculate a weighted average of the accumulated disturbance intensity for each species, using each species’ relative occurrence across the different disturbance treatments at the end of the experiment as weights. Next, we investigate the effects of the disturbance treatments on taxonomic and functional turnover in community composition (rate of change) and on changes in the spatial heterogeneity of taxonomic and functional compositions within treatments between the initial situation (T0) and each subsequent sampling moment. Finally, we investigate the effects of the disturbance treatments on changes in various diversity and functional indices between the initial situation (T0) and each subsequent sampling moment.

#### Disturbance sensitivity of species

We calculated weighted averages of species abundances over the experimental disturbance gradient to investigate whether certain species are associated with no, low, or higher disturbance treatments. Here, each treatment was assigned a value based on the annual cumulative disturbance intensity for each treatment (i.e., C =0, I_low_F_high_ = 3, I_high_F_low_ = 3, and I_high_F_high_= 9), reflecting the annual Swept Area Ratio (SAR) used to express fishery impact. Weighted averages were then determined for all species found in at least three samples for T3, T6, and T7. For this specific analysis, we excluded T0 as this was pre-disturbance, and T1 as this was only sampled one week after the initial disturbance was applied. The mean and standard errors were then determined from the weighted average over the three time periods. In addition to the weighted average, we also determined the proportions of each species across the different treatment types.

#### Community turnover

We first assessed the similarity in community composition between treatments within locations in the pre-disturbed state (T0) using a redundancy analysis with a treatment-by-location interaction as a constraining variable. This was done using an Euclidean distance method on Hellinger-transformed abundances for the taxonomic community, and on the community-weighted mean for the functional community. Model significance was tested using the *‘adonis2’* function from the ‘vegan’ package (Oksanen et al., 2024). A post-hoc test was employed in the case of a significant model by sequentially testing group differences between treatments per location using the ‘*pairwise.adonis2’* function from the ‘pairwiseAdonis’ package (Martinez Arbizu, 2017). The Hellinger standardisation (square root of the proportion of a species in a sample over all individuals) is a standardisation step that simultaneously helps minimise the effects of vastly different sample total abundances, as typically found in benthos samples (i.e., which may sometimes contain only a few big bivalves but in other samples hundreds of small worms). Moreover, the Hellinger standardisation allows for the application of linear methods to community data (Legendre & Gallagher, 2001). P-values were corrected for multiple tests using a Bonferroni correction (Dunn, 1961). Here, only treatment differences within time periods (T’s) are tested, not between time periods. Because the treatments significantly differed in terms of community and functional composition in most locations at T0, reflecting substantial spatial heterogeneity (Supplementary Figures S2, S3), we further investigated community changes by comparing changes over time for each treatment starting from the pretreatment sampling, instead of simply comparing community composition to the control treatment at any time, as this would not account for initially existing differences among treatments. We calculated the Euclidean distance for each sampling point compared to T0 at the same location using the Hellinger-transformed community data and community-weighted functional trait composition to investigate the total shift since the start of the experiment. Moreover, to investigate the spatial heterogeneity of the communities (community dissimilarity) within each experimental plot (treatment-location combination), we also calculated Hellinger and Euclidean distances for the taxonomic and trait communities within each combination of location, treatment, and time period. To analyse differences in means of the Hellinger and Euclidean distances across treatments over time, we employed linear mixed models with time period and treatment as interactive effects, distance measures as the response variable, and a random effect of sampling station nested within location. Models were then bootstrapped with 10.000 parametric iterations using the *‘bootstrap_model’* function from the “parameters” R package (Lüdecke et al., 2020). Confidence intervals were calculated, and contrast comparisons on the bootstrapped estimates were then conducted using the *‘emmeans’* function from the “emmeans” R package (Lenth, 2024) and the ‘*model_parameters’* function from the “parameters” (Lüdecke et al., 2020) R package with Sidak adjustment (Sidak, 1967) for the p-values.

#### Diversity and functionality

For each sampling station, alpha-diversity was calculated using Hill numbers to provide a unified, interpretable framework for diversity assessment (Hill 1973). Hill numbers present diversity as the effective number of species, allowing direct and meaningful comparisons between assemblages and over time (Chao et al., 2014). Within the Hill number framework, parameter *q* determines the sensitivity of the metric to the dominance of species (Hill 1973, Chao et al., 2014). With Hill *q* = 0, equal weight is given to both rare and common species, effectively calculating species richness. Hill *q* = 1 (hereafter referred to as ENS1) and Hill *q* = 2 (hereafter referred to as ENS2) return the effective number of species derived from Shannon and Simpson indices. Evenness was calculated as the ratio between the value of ENS1 and ENS2 (Hill 1973; Ricotta and Avena, 2002). To examine changes from a functional trait perspective, we also determined the functional diversity of each sampling station using Rao’s quadratic entropy (hereafter Rao’s Q) (Botta-Dukát, 2005) using the ‘*dbFD*’ function from the ‘FD’ package (Laliberté et al., 2014). Moreover, we examined changes in the proportion of long-lived species, suspension feeders, and the combined proportion of predators and scavengers. For each metric, we calculated the log change ratio between T0 and each subsequent sampling period for all variables using the natural log and by adding a value of one to each value to avoid dividing by zero or taking the natural log of zero. We then used linear mixed models with sampling stations nested in location as a random effect and an interaction between treatment and time period to compare mean log change ratios between treatments over time since T0. The same bootstrapping method was then applied to these models as described above.

## Results

### Disturbance sensitivity of species

We calculated weighted averages of species abundances along the experimental disturbance gradient to identify species that are sensitive to disturbance versus those that are tolerant. The lowest weighted average Swept Area Ratio (SAR) was found for the polychaete worm *Sthenelais boa*, which only occurred in the Control treatment (Figure 2). Other species, such as the polychaete worms *Travisia forbesii*, the bivalves *Barnea candida* and *Magallana gigas*, and Actiniaria (anemones), had a low weighted average SAR (Figure 2A) and were also primarily found in the Control treatment but also occasionally in the low intensity and high frequency (I_low_F_high)_, and high intensity and low frequency (I_high_F_low)_ treatments (Figure 2B). No species were clearly associated with high intensity and high frequency (I_high_F_high_) treatments based on the proportional distribution of species over the treatments. Species associated with high disturbance intensity (high SAR values), such as those found for the polychaete worms *Scolelepis* sp. and *Vermiliopsis striaticeps*, and the gastropod *Retusa obtusa* (Figure 2A), proportionally occur in all treatments (Figure 2B).

**Figure 2.**
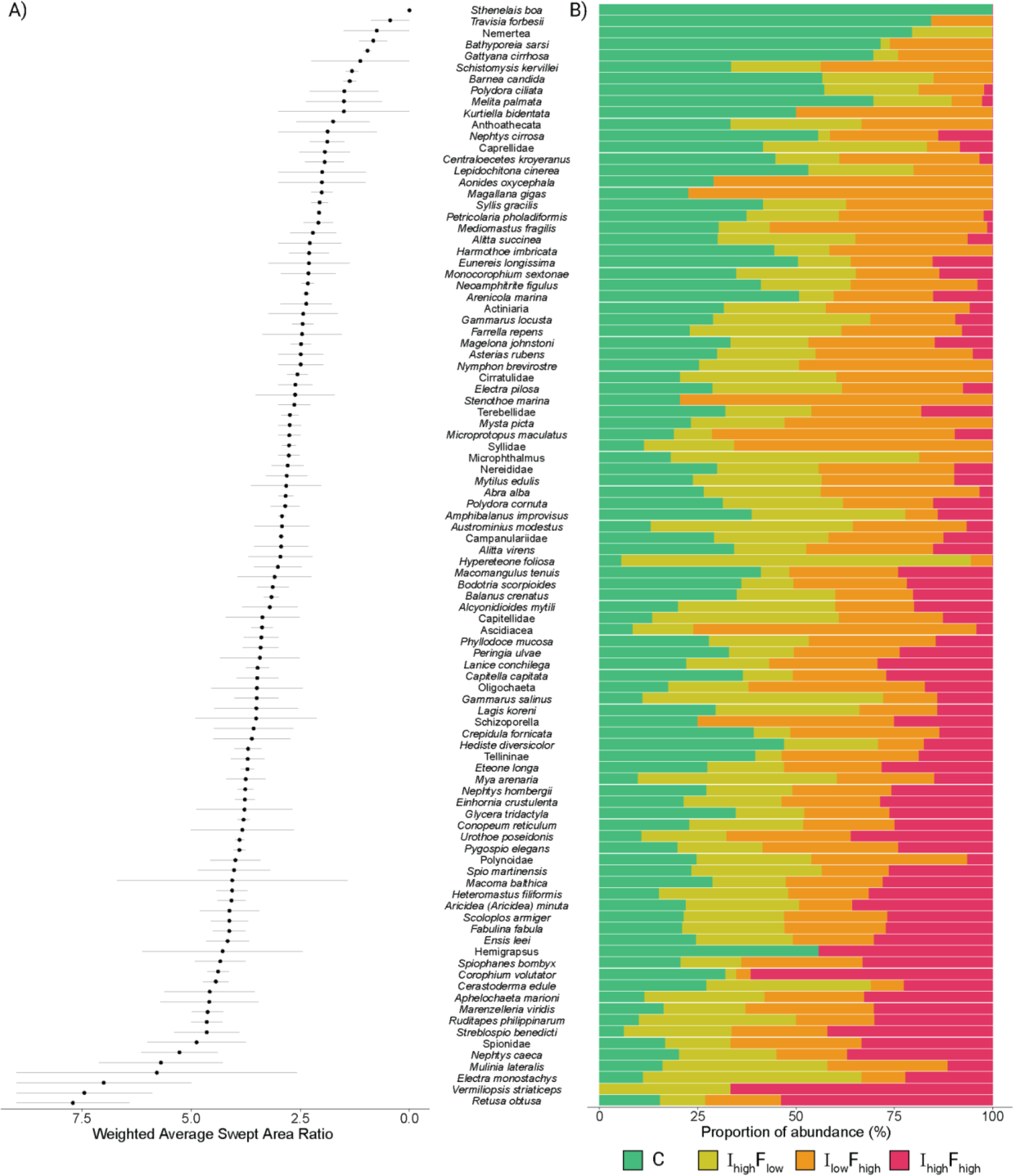
A) average disturbance frequency at which each species was observed (ranked from low to high) calculated as a weighted average Swept Area Ratio, using the observed abundances in the different disturbance treatments. Means and standard errors are calculated over sampling periods T3, T6, and T7. High values at the bottom reflect species that occur in plots with highly disturbed treatments, and low values at the top indicate species that occur in plots with more stable conditions. B) Relative proportion of species abundances distributed over the different treatments over T3, T6, and, T7. C is Control, I_high_F_low_ is High intensity, Low frequency, I_low_F_high_ is Low intensity, High frequency, and I_high_F_high_ is High intensity, High frequency.

### Changes in community structure over time

We observed strong spatial heterogeneity within the study sites, as treatment plots within sites already showed differences in both species and trait communities before the first experimental disturbance. This strong spatial heterogeneity between assigned treatment plots could not be accounted for by the randomised placement of replicate treatments (Supplementary Figures S2 and S3). These initial differences in species and trait communities pre-disturbance between the different treatments complicated the direct conclusions about disturbance effects on community composition, and comparing means between treatments using a standard randomised block design. To overcome this limitation, we investigated the community turnover rate and dissimilarity, as we expect turnover rate to increase with disturbance due to increasing changes in abundance and species identity (Kaarlejärvi et al., 2021). Throughout the study period, all communities, including the controls, shifted strongly in composition, indicating large seasonal variations in species and traits over time (Figure 3A, B) but also showing differences between treatments in their degree of turnover. Turnover of the species and trait communities was largest in the most frequently disturbed I_high_F_high_ treatments and differed significantly from the Control treatment (Figure 3A, B; Supplementary Table S3). The turnover in the intermediate I_low_F_high_ and I_high_H_low_ treatments for both the species and trait community was between the turnover rate for the Control and I_high_F_high_ treatments, although they were not significantly different throughout the entire experimental period (Figure 3A, B; Supplementary Table S3).

**Figure 3.**
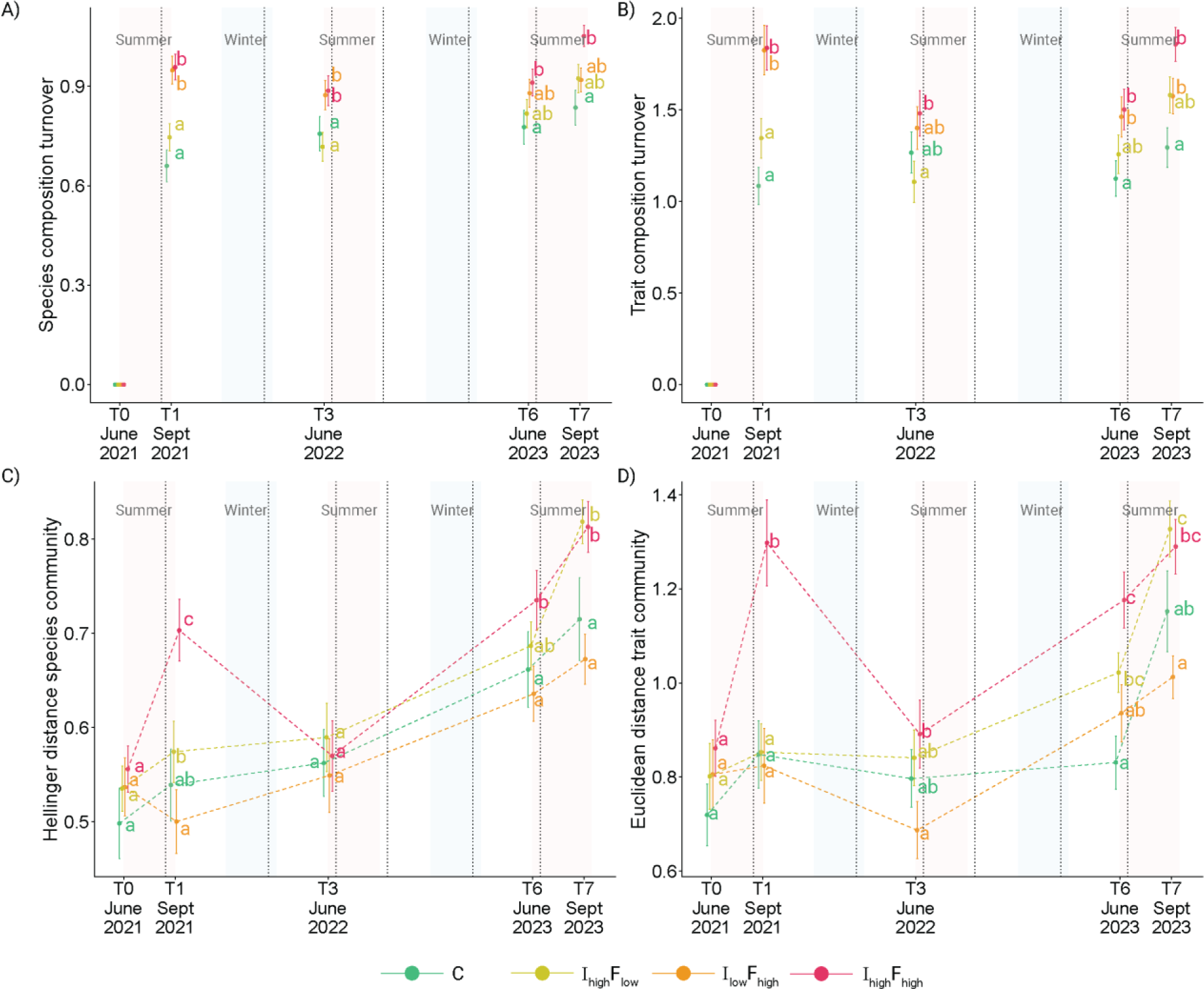
Log change ratio’s over time reflecting community turnover compared to T0 for A) Hellinger distance of the species community and B) Euclidean distance of the functional composition, and the community dissimilarity throughout time for C) Hellinger distance of the species community and D) Euclidean distance of the trait community. The mean turnover with standard error is shown for each treatment per time period. Dashed vertical grey lines represent disturbance events. Background tiles indicate seasons. Significant differences in means between treatments within time periods are indicated by a compact letter display. C is Control, I_high_F_low_ is High intensity, Low frequency, I_low_F_high_ is Low intensity, High frequency, and I_high_F_high_ is High intensity, High frequency.

Species and trait community dissimilarity (heterogeneity) were not different between treatments at the start of the experiment, pre-disturbance, but then diverged between treatments (Figure 3C-D). Immediately after the first disturbance, the I_high_F_high_ treatment had a significantly higher within-treatment dissimilarity (mean ± SE = 0.70 ± 0.033 for species and 1.29 ± 0.090 for traits) than the other treatments (Figure 3C-D, Supplementary Table S3). The I_high_F_low_ treatment (mean ± SE = 0.57 ± 0.033 for species and 0.85 ± 0.060 for traits), which was equally intensely disturbed during the first disturbance event, had a higher mean dissimilarity than the Control (mean ± SE = 0.54 ± 0.038 for species and 0.85 ± 0.071 for traits) and I_low_F_high_ (mean ± SE = 0.50 ± 0.034 for species and 0.81 ± 0.078 for traits) treatments, but this was only significant in the comparison to I_low_F_high_. As the experiment continued, the within-treatment dissimilarity of the species and trait communities increased over time for all treatments. Still, the I_high_F_low_ and I_high_F_high_ treatments diverged from the I_low_F_high_ and Control treatments with significantly greater dissimilarity at the end of the experiment for the species community (I_high_F_low_ = 0.82 ± 0.023 and 1.32 ± 0.060, I_high_F_high_ = 0.81 ± 0.030 and 1.27 ± 0.055, I_low_F_high_ = 0.67 ± 0.027 and 1.00 ± 0.045, Control = 0.72 ± 0.044 and 1.15 ± 0.086, for species and traits, respectively) (Figure 3C; Supplementary Table S3).

### Diversity and functionality

We found no significant differences in the change in number of species during the experimental period between the treatments (Figure 4A; Supplementary Table S3). A strong decrease was observed in species richness in T1 compared to T0 though this was similar between treatments, and returned to original values over the course of the experiment (Figure 4A; Supplementary Table S3). Abundance weighted metrics of diversity (i.e., ENS1, ENS2, and evenness) did show some statistically significant differences in change between treatments, however, these did not reveal a single clear direction of change (Figure 4A; Supplementary Table S3). Contrastingly, functional diversity (Rao’s Q) did show a uniform response in the change in diversity over time between the treatments (Figure 5A). The functional diversity in the I_high_F_high_ and I_low_F_high_ treatments both decreased in all time periods compared to T0, whereas it increased in the C treatment for all time periods compared to T0 and was highly variable but overall stable for the I_high_F_low_ treatments in all time periods compared to T0 (Figure 5A) (Figure 5A, Supplementary Table S3). We also found specific shifts in functional groups over time. The proportion of long-living species at the end of the experiment was higher than at T0 for all treatments (Figure 5B). However, this increase was only visible in the last time period (T7) for the high-frequency treatments (I_high_F_high_ mean ± SE = 0.050 ± 0.031 and I_low_F_high_ mean ± SE = 0.041 ± 0.024) and was significantly lower than for the Control (mean ± SE = 0.11 ± 0.021) and I_high_F_low_ (mean ± SE = 0.10 ± 0.022) treatments. Moreover, the change in long-living species declined strongly in the I_high_F_high_ treatment at T3 (mean ± SE = -0.055 ± 0.024) (Figure 5B, Supplementary Table S3). Similarly, the high-frequency treatments (I_high_F_high_ and I_low_F_high_) were associated with higher increases in scavengers and predator species and lower increases in suspension feeders compared to the low-frequency disturbance (I_high_F_low_) and controls (Figure 5C, D, Supplementary Table S3).

**Figure 4.**
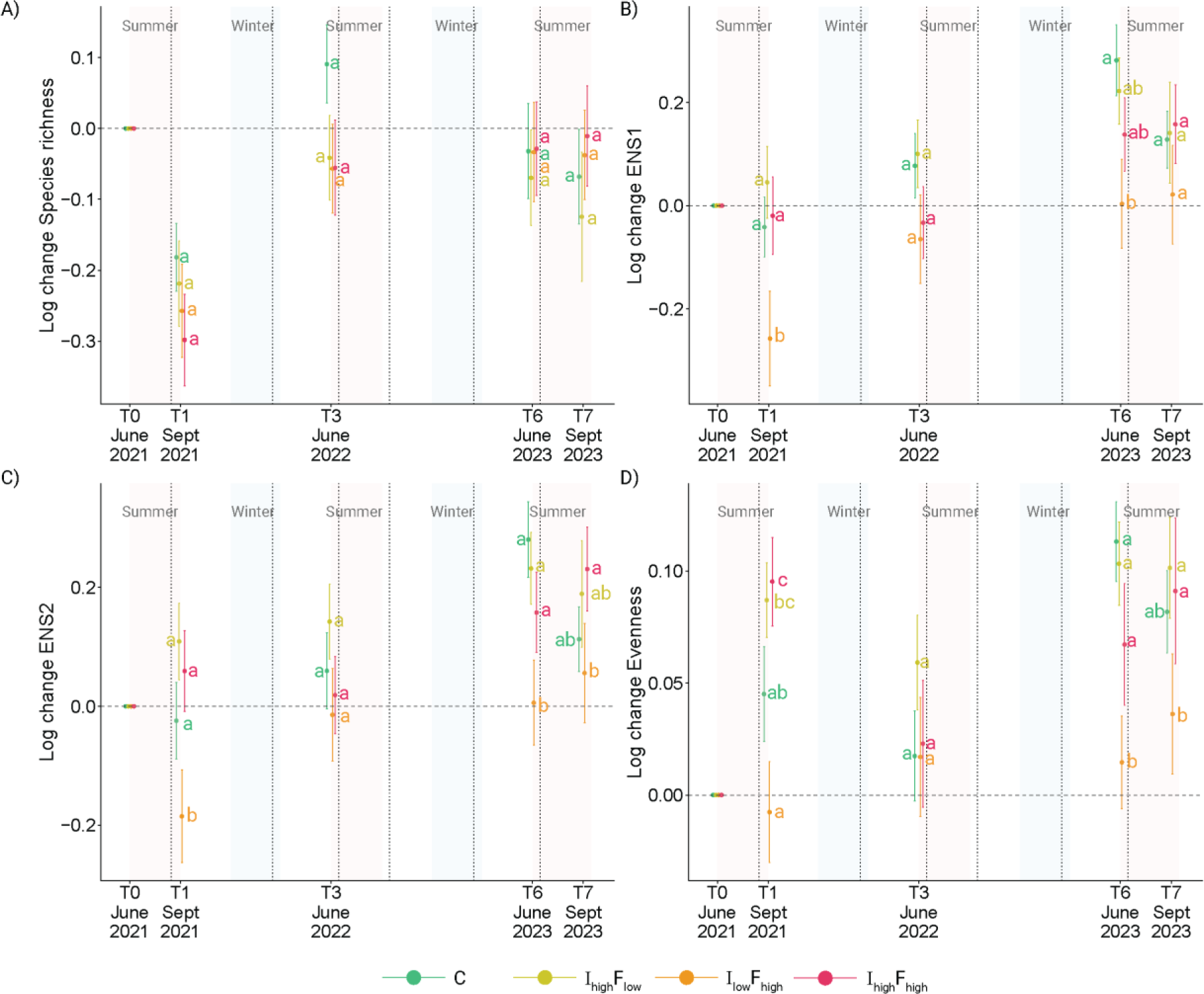
Log change ratios over time compared to T0 in A) species richness, B) ENS1, C) ENS2, and D) Evenness (ENS2/species richness). The mean with standard error is shown for each treatment per time period. Vertical dashed grey lines represent disturbance events. Background tiles indicate seasons. Significant differences in means between treatments within time periods are indicated by a compact letter display. C is Control, I_high_F_low_ is High intensity, Low frequency, I_low_F_high_ is Low intensity, High frequency, and I_high_F_high_ is High intensity, High frequency.

**Figure 5.**
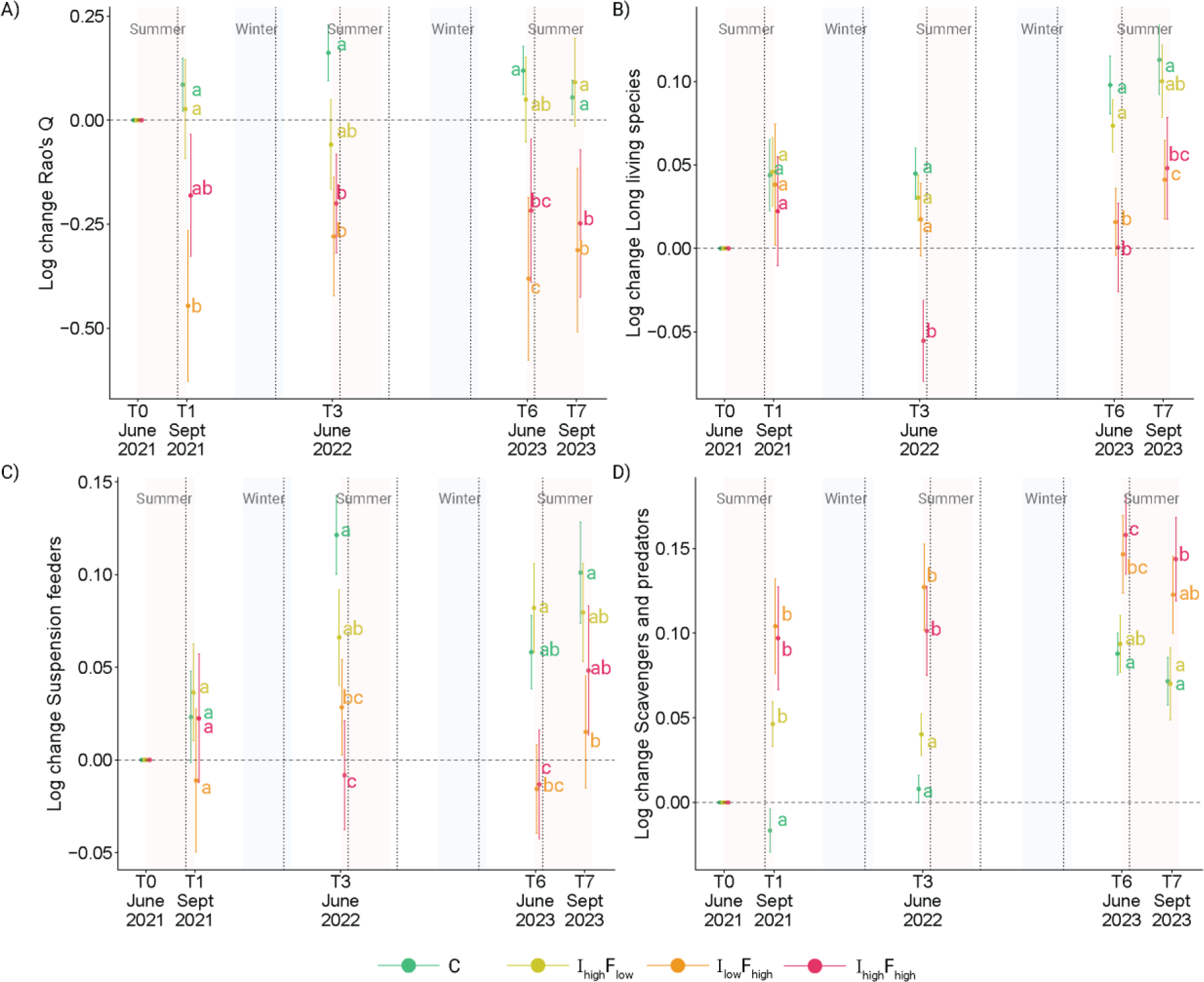
Log change ratio’s over time compared to T0 for A) Rao’s Q, B) proportion of long-living species (>6 years), C) proportion of suspension feeders, and D) proportion of scavengers and predators. The mean with standard error is shown for each treatment per time period. Vertical dashed grey lines represent disturbance events. Background tiles indicate seasons. Significant differences in means between treatments within time periods are indicated by a compact letter display. C is Control, I_high_F_low_ is High intensity, Low frequency, I_low_F_high_ is Low intensity, High frequency, and I_high_F_high_ is High intensity, High frequency.

## Discussion

Marine Protected Areas are widely recognised as key tools for conserving marine ecosystems and mitigating the impacts of anthropogenic pressures (Lubchenco & Grorud-Colvert, 2015; Grorud-Colvert et al., 2021). However, their effectiveness in dynamic soft-bottom systems remains poorly understood. These systems are characterised by high levels of physical stress through strong hydrodynamic forces to which benthic communities are argued to be inherently adapted. This raises the question of whether protection from additional anthropogenic disturbance provides ecological benefits in environments where natural forces already play a dominant role in structuring communities. We use the Dutch Wadden Sea as a model system to evaluate the effects of bottom disturbance within Marine Protected Areas and measure ecological responses to reintroduced human impact. Our findings demonstrate that even low-intensity mechanical disturbance from relatively light trawling gear significantly affects macrozoobenthic community structure and functional composition. Though standard diversity metrics, i.e., species diversity metrics following the Hill number framework, were not affected, we observed impacts across multiple other community metrics, including shifts in functional groups, declines in functional diversity and increased turonover of species communities. This suggests that anthropogenic disturbance remains a relevant ecological pressure, even in systems that may appear resilient due to their dynamic nature.

### Community effects

Our results show that specific species are associated with low levels of disturbance, underscoring the role of Marine Protected Areas in safeguarding sensitive benthic communities. The sensitivity of species to bottom disturbance is typically associated with specific life history traits (Kindsvater et al., 2016). For example, epibenthic, large, and sessile species are often found in areas of the North Sea that are less or not disturbed by bottom trawling (van Denderen et al., 2015). In contrast, highly trawled areas often contain more mobile, deep-living, and smaller species that are less susceptible to passing nets (van Denderen et al., 2015). Similarly, we find sessile, epibenthic species, such as Actiniaria species (anemones), to be associated with low levels of disturbance. Conversely, no clear set of species was consistently associated with high-disturbance treatments. Tolerant species were found across all treatments, including the control treatments. This pattern indicates that disturbance primarily affects species relative abundances rather than presence/absence, and highlights the potential of Marine Protected Areas to support populations of disturbance-sensitive species.

Community composition showed high variability and lacked a clear directional shift, likely due to pre-existing differences between treatment plots at the start of the experiment. Control treatments already differed pre-disturbance in taxonomic and functional composition from the other treatments in most locations, even at the relatively small scale of the experiment. To overcome this, we instead examined community turnover per location since the start of the experiment. This turnover was high across all treatment plots, including the control. This may reflect the general finding of other studies that substantial unexplained variation remains when assessing the effects of bottom disturbance on macrobenthic communities (Sköld et al., 2018). Community assembly in highly dynamic coastal areas, like the Wadden Sea, is hypothesised to be primarily driven by random processes (Sloots et al., 2025). Indeed, this is reflected in the high community turnover of species and trait communities, as well as in the strong heterogeneity within treatment locations. Still, the turnover was more substantial in plots that were more frequently disturbed, whereas heterogeneity increased with higher-intensity plots. Increased community turnover is commonly found following disturbance (e.g., Kaarlejärvi et al., 2021; Swenson et al., 2012) due to greater changes in species identity and abundance than in stable non-disturbed systems (Sousa, 1984; Supp & Ernest, 2014). Then, the long and ubiquitous presence of human bottom disturbance will have contributed to the system’s high stochasticity of community assembly. This presents a challenge for conservation efforts, as increased unpredictability complicates the monitoring and evaluation of targeted conservation measures, and protected areas must be large enough to capture the scale and heterogeneity of community assembly processes and effectively protect all biodiversity (Economo, 2011).

### Diversity and functionality

While standard diversity measures, such as species richness or the effective number of species, remain valuable for quantifying change in biodiversity, they inherently fail to account for species traits and functions (Chao et al., 2014; Hillebrand et al. 2018; Eriksson and Hillebrand 2019). This can obscure meaningful ecological shifts, especially in dynamic systems where community composition is highly variable and disturbance may substitute one set of species for another without affecting taxonomic diversity indices. Indeed, none of our diversity indices show consistent unidirectional responses to disturbance, and changes are largely similar between treatments over time, despite using multiple diversity indices with variable weighing of species dominance. Contrastingly, using functional diversity, clear responses to the disturbance treatments are visible. Rao’s Q demonstrates a significant decline with more frequent disturbances, regardless of their intensity, suggesting a reduction in functional diversity due to bottom disturbance in which the frequency at which a disturbance occurs is more important than its immediate strength. These findings are consistent with McLaverty et al. (2024), who report insignificant decreases in species diversity with more frequent disturbances by shrimp trawling and significant declines in functional diversity. Functional diversity is thus more sensitive and should be prioritised in impact assessment, as changes in functional composition caused by bottom trawling may affect ecosystem stability and functions (Hillebrand et al., 2008).

Considering changes in terms of functional groups provides a more nuanced picture of community responses to disturbance than taxonomic diversity metrics alone. We observed clear changes in the relative abundance of functional groups depending on the frequency and intensity of bottom disturbance. Suspension feeders increased at the control and low-frequency sites, but not at the high disturbance frequency site. Contrastingly, scavengers and predators increased in all disturbed sites, likely due to greater food availability following the direct mortality of organisms caused by passing trawls, a finding well-documented in bottom-trawling studies (Tillin et al., 2006; McLaverty et al., 2024) as well as studies on the impacts of anchoring (Broad et al., 2020; Davis et al., 2025). This increase in scavengers and predators was clear within a week (at T1) after the first disturbance event at all disturbed sites. However, in the low-frequency treatment, the community composition recovered to control levels shortly thereafter. Nevertheless, by the end of the experiment, both the control and low-frequency treatments showed an increased proportion of scavengers and predators. However, the magnitude of change was significantly lower compared to high-frequency sites. This can either be the cause of a population-wide increase in scavenger-predators within the system, or because the effects of bottom disturbance extend beyond the directly impacted area. In contrast, the limited decrease in suspension feeders was observed only at the frequently disturbed sites. Suspension feeders are generally sessile and particularly sensitive to disturbance, especially due to the negative effects of increased suspended sediment (Thrush et al., 2021). This decline was not evident in the first sampling immediately following the initial disturbance, likely due to delayed responses (Tilman et al., 1994), but similar declines in suspension feeders have been reported in other studies (Tillin et al., 2006; de Juan et al., 2007; Oberle et al., 2016). We also found that long-lived species increased relative to the high-frequency sites in both the control and low-frequency disturbance treatments. Longevity is a well-established proxy for sensitivity to disturbance, as it is associated with traits like delayed reproduction and lower egg survival, which slow population recovery (Rijnsdorp et al., 2018). Overall, frequent bottom disturbance alters the longevity and feeding group composition of benthic communities, leading to shifts across different spatial scales. These shifts may have significant implications for food web dynamics and the recovery potential of populations, particularly with the loss of species that have slower life-history strategies (Tillin et al., 2006; Rijnsdorp et al., 2018; van de Wolfshaar et al., 2020). Marine Protected Area design should then consider the scales at which recovery and disturbance operate, especially given potential edge effects that operate at different functional levels.

### Time lags of Marine Protected Areas and disturbance effects

The experimental sites, although protected, likely do not represent a completely undisturbed community. We conducted the experiment in previously identified Marine Protected Areas (Meijer et al., 2025), which were established only six years before the start of our experiments in relatively disturbed areas, to allow for at least some potential initial recovery of disturbance-sensitive species. Proper Marine Protected Areas, apart from the 2005 reference area in tidal basin ‘Schild’, did not exist in the Dutch Wadden Sea until 2015. Only then, through agreements among the fishery sectors, nature organisations, northern provinces, and the Ministry of Agriculture, were areas protected from bottom trawling for mussel seed and shrimp fishery (PRW, 2014). This left only six years between the initial closure of the areas to bottom disturbance and the start of the experiment. Monitoring programs initiated to track developments in these areas have, so far, also not observed clear, persistent differences between them and non-protected areas in the first six years after closure. However, these programs also lack proper pre-closure measurements, which complicates accurate assessment (Craeymeersch et al., 2022). Moreover, subtle changes in the macrozoobenthic community in the reference area Schild was only measurable after ten years (Glorius et al., 2018). Thus, the control and treatment plots we used in our experiment are likely still developing and recovering from historical disturbance regimes, which diminishes the possibility of detecting differences between treatments. Moreover, the strong and long history of human disturbances in the system may have degraded the entire meta-community of the wider system (Thrush et al., 2021a), apparent from the large scale changes in species since human settlement (Lotze et al., 2005), leaving the system without undisturbed communities and, therefore, with an impoverished regional species pool, which complicates local recovery (Lundquist et al., 2006; Thrush et al., 2008). The experimental trawling intensities used in our experiment (SAR’s of 3 and 9) are within the range commonly found within the system, though higher SAR values (SAR > 15) can be found (ICES, 2021), implicating even stronger realised impacts throughout the system. Importantly, the cumulative effects of repeated disturbances became more pronounced over time, suggesting that short-term or single-impact studies may substantially underestimate the actual ecological consequences of bottom disturbance.

### Conclusion and Management Implications

In this large-scale experimental study, we show that even trawling with light gear can significantly affect both the taxonomic and functional dynamics of benthic communities in highly dynamic coastal systems. The observed shifts toward scavenging species, combined with declines in long-lived and suspension-feeding organisms, suggest long-term alterations to the ecosystem with potential consequences for its resilience. The effects observed, therefore, conflict with the Natura 2000 regulations in effect in the study area. Especially given that our experiment solely focused on the physical disturbance caused by the trawl, without considering the additional trophic effects of biomass removal, the realised effects of this bottom trawling are likely to be even more impactful. Considering this, it is likely that the long history of shrimp fishing in the Wadden Sea has influenced the current status of macrozoobenthic communities in the ecosystem. Given its Natura 2000 designation, the World Heritage Status, and ecological importance, restricting bottom-disturbing activities through Marine Protected Areas is crucial for achieving conservation goals.

Our findings further highlight that Marine Protected Areas play a key role in safeguarding ecological integrity by protecting sensitive and functionally important species. Even in environments with high natural variability, anthropogenic disturbance remains an additional stressor that alters community composition and ecosystem function. Standard diversity metrics fail to capture the full impact of anthropogenic pressures on ecosystems and the ecological benefits provided by Marine Protected Areas. Functional diversity metrics and community metrics offer complementary perspectives by incorporating ecological roles and species identity, distinguishing between changes in species identity and true functional losses. The observed changes in long-lived and suspension-feeding organisms within Marine Protected Areas compared to impacted areas underscores the importance of including functional traits in impact assessment. This is especially important in management contexts where conservation aims should extend beyond maintaining species numbers to safeguarding functional integrity and stability of communities in the face of anthropogenic pressure, for which Marine Protected areas are thus a key conservation tool.

## Acknowledgements

The authors would like to thank the crew on the NIOZ Research Vessel “Navicula”, in particular: Bram Fey, Hein de Vries, Klaas-Jan Daalder, and Willem-Anton Schagen, and everyone who was involved in the sampling and the processing of samples in the laboratory. In particular, we would like to thank: Aldara Gil Garcia, Amin Niamir, Andy Niele, Annemieke van Schie, Bartel Komin, Bianka Rasch, Bob Monnich, Daniel Varley, Daphne Somers, Demi Damstra, Dirk Schagen, Dorien Oude Luttikhuis, Evaline van Weerlee, Felianne van Leersum, Franka Lotze, Isa Faber, Jeroen Kooijman, Julia Dorigo, Jynthe Plakman, Juan Schiaffi, Katrin Rehlmeyer, Konstantina Bairaktari, Lisa Mijnheer, Livia Brunner, Loran Kleine Schaars, Maureen Sikkema, Marta Ferraro, Martin Mensen, Nadia Hijner, Puck Heerink, Reyhaneh Roohi, Roos van Dorp, Rozemijn van Dam, Rueben Millenaar, Tom Dragt, and Yannick Hill. In addition, we would like to thank Leo Visser and Diederik Visser of the fishing vessel LO8 (Trijntje) for performing the experimental disturbances for this study and Ingrid Tulp for her critical views on earlier versions of the manuscript. This study is part of project “Waddentools: habitatheterogeniteit” (also known as “Waddenmozaïek”), registered under reference number WF2018-187059, and funded by the Waddenfonds, the Directorate General for Public Works and Water Management (Rijkswaterstaat) and the provinces of Noord-Holland, Fryslân and Groningen.

## Data Availability

The codes and data produced for this manuscript will be available in the dataverseNL (DANS) repository upon publication.

## Declaration of competing interests

The authors declare that they have no known competing financial interests or personal relationships that could have appeared to influence the work reported in this paper.

## Author contributions

**Kasper J. Meijer**: Conceptualization, methodology, data curation, formal analysis, investigation, writing - original draft, writing - review and editing

**Oscar Franken**: Conceptualization, methodology, writing - review and editing

**Laura L. Govers**: Conceptualization, methodology, writing - review and editing

**Tjisse van der Heide**: Conceptualization, methodology, writing - review and editing

**Eva Willebrands**: Investigation, writing - review and editing

**Mark E. de Wilt**: Investigation, writing - review and editing

**Bas de Wit**: Investigation, writing - review and editing

**Han Olff**: Conceptualization, methodology, supervision, writing - review and editing

## Notes

### Competing Interest Statement

The authors have declared no competing interest.

