## Supplementary material for "Experimental Disturbance-Induced Shifts in Benthic Functional Diversity: the importance of marine protected areas in soft-bottom ecosystems"

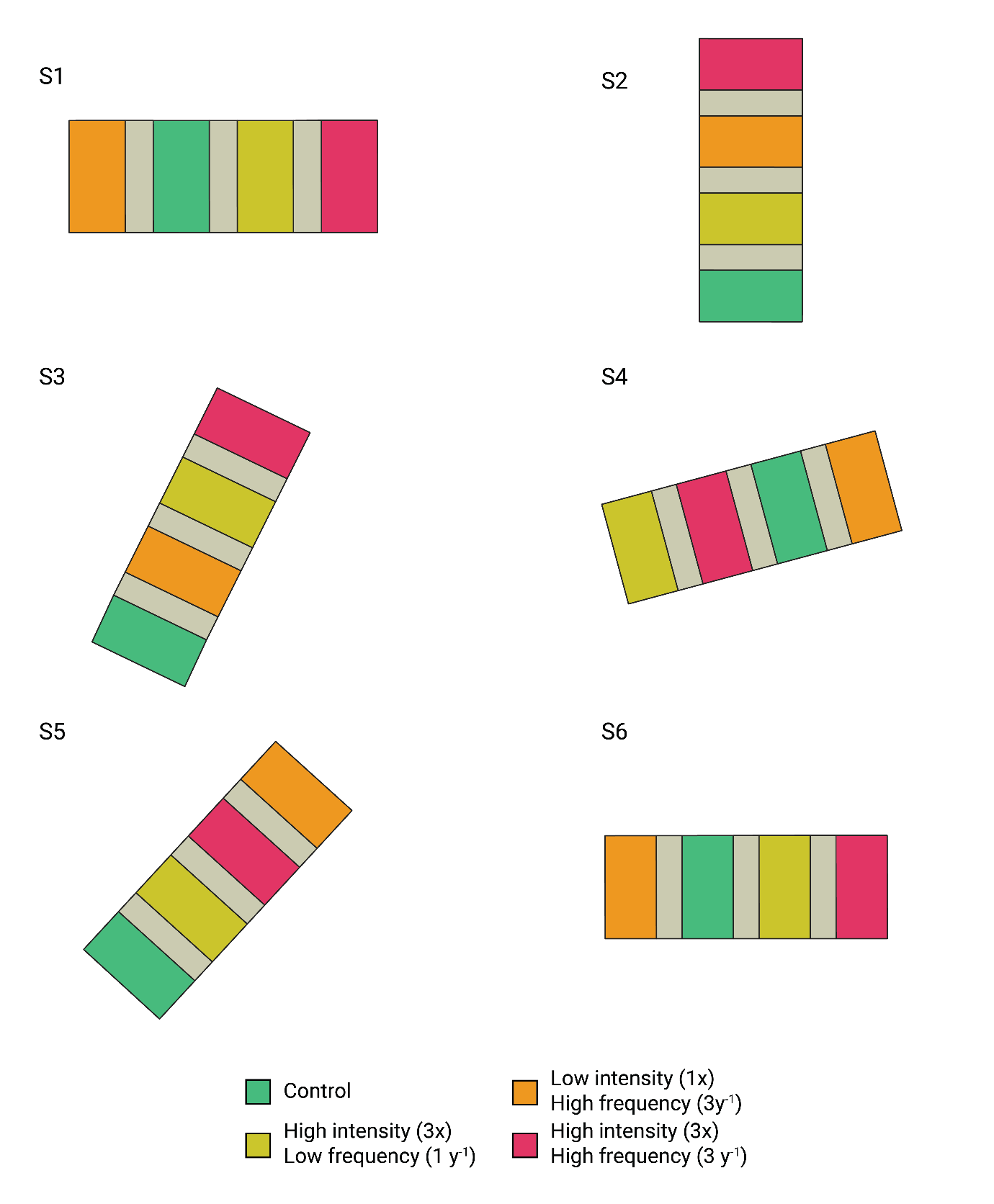
**Supplementary Figure S1.** Close-up of experimental sites with the randomised treatment order. For site location and orientation refer to Figure 1.

##### **Supplementary Table S1.** Overview of the number of included samples in the analysis per time period, location and treatment.

| Sampling Period | Treatment | Number of Samples |
| --- | --- | --- |
| T0 | Control | 52 |
| T0 | I_high_F_low_ | 51 |
| T0 | I_low_F_high_ | 47 |
| T0 | I_high_F_high_ | 54 |
| T1 | Control | 52 |
| T1 | I_high_F_low_ | 52 |
| T1 | I_low_F_high_ | 51 |
| T1 | I_high_F_high_ | 49 |
| T3 | Control | 54 |
| T3 | I_high_F_low_ | 54 |
| T3 | I_low_F_high_ | 53 |
| T3 | I_high_F_high_ | 54 |
| T6 | Control | 54 |
| T6 | I_high_F_low_ | 54 |
| T6 | I_low_F_high_ | 54 |
| T6 | I_high_F_high_ | 54 |
| T7 | Control | 53 |
| T7 | I_high_F_low_ | 51 |
| T7 | I_low_F_high_ | 54 |
| T7 | I_high_F_high_ | 53 |

##### **Supplementary Table S2.** Overview of traits and modalities. Adapted from Meijer et al., 2023.

| **Trait** | **Modality** | **Description** |
| --- | --- | --- |
| Bioturbation type | Epifauna | No bioturbating activity due to epifaunal living environment |
|  | Surficial modifier | Any invertebrate whose sediment reworking activity is restricted to the uppermost (~1 cm) sediment layers (Solan et al 2004) |
|  | Upward conveyor | Head-down oriented fauna which causes active sediment movement from depth to the surface (François et al. 1997) |
|  | Downward conveyor | Head-up oriented fauna which causes active sediment movement from the surface to depth through their gut (François et al. 1997) |
|  | Biodiffuser | Fauna that randomly moves sediment over short distances causing diffusive mixing (François et al. 1997) |
|  | Regenerator | An invertebrate that transfers sediment at depth to the surface where it is washed away and replaced by the sediment of surficial signature (Gardner et al., 1987) |
| Adult living depth (cm) | surface | Surface living species |
|  | >0 and ≤ 3 | Species living between 0-3 cm depth in the sediment |
|  | >3 and ≤ 8 | Species living between 3-8 cm depth in the sediment |
|  | >8 and ≤ 15 | Species living between 8-15 cm depth in the sediment |
|  | >15 and ≤ 25 | Species living between 15-25 cm depth in the sediment |
|  | >25 | Species living deeper than 25 cm in the sediment |
| Adult body size (mm) | ≤ 5 | Body size smaller then 5 mm |
|  | >5 and ≤ 10 | Body size between 5-10 mm |
|  | >10 and ≤ 20 | Body size between 10-20 mm |
|  | >20 and ≤ 40 | Body size between 20-40 mm |
|  | >40 and ≤ 80 | Body size between 40-80 mm |
|  | >80 and ≤ 160 | Body size between 80-160 mm |
|  | >160 | Body size larger than 160 mm |
| Feeding Mode | Deposit-feeder | Taxa that forage along the surface and ingest soft parts of the sediment and so digesting and assimilating organic matter |
|  | Suspension feeder | Taxa that obtain food by filtering particles from the water column |
|  | Grazer | Taxa that graze on plants and algae |
|  | Opportunist/scavenger | Opportunistically feeding taxa |
|  | Predator | Predatory taxa |
| Longevity (y) | ≤1 | Taxa that live less than 1 year |
|  | >1 and ≤ 3 | Taxa that live between 1-3 years |
|  | >3 and ≤ 6 | Taxa that live between 3-6 years |
|  | >6 and ≤ 10 | Taxa that live between 6-10 years |
|  | > 10 | Taxa that live longer than 10 years |
| Age of sexual maturation (y) | ≤1 | Sexually mature within 1 year |
|  | >1 and ≤ 2 | Sexually mature between 1-2 years |
|  | >2 and ≤ 5 | Sexually mature between 2-5 years |
|  | >5 and ≤ 10 | Sexually mature between 5-10 years |
|  | >10 | Sexually mature after 10 years |
| Reproductive frequency | Continuous / >= 2x per year | Two or more reproductive events per year |
|  | Annual 1x | One reproductive event per year |
|  | Biennial | Reproduces every other year |
|  | Semelparous | Reproduces once in a lifetime |
| Fecundity | ≥1 and ≤ 50 | Reproductive output of 1-50 offspring over its lifetime |
|  | >50 and ≤ 500 | Reproductive output of 50-500 offspring over its lifetime |
|  | >500 and ≤ 2.500 | Reproductive output of 500-2.500 offspring over its lifetime |
|  | >2.500 and ≤ 10.000 | Reproductive output of 2.500-10.000 offspring over its lifetime |
|  | >10.000 and ≤ 20.000 | Reproductive output of 10.000-20.000 offspring over its lifetime |
|  | >20.000 and ≤ 100.000 | Reproductive output of 20.000-100.000 offspring over its lifetime |
|  | >100.000 | Reproductive output of more than 100.000 offspring over its lifetime |
| Mobility | Sessile | Taxa that are sessile and do not move |
|  | Swim/float | Taxa that can move around by swimming or floating through the water column |
|  | Crawl/walk | Taxa that move around by crawling or walking |
|  | Burrow/tube | Taxa that move around by burrowing through the sediment or by creating tubes |
| Adult living habitat | Tube | Taxa that live in tubes |
|  | Burrow | Taxa that burrow through the sediment |
|  | Free-living | Taxa that live freely |
|  | Crevice | Taxa that live in between crevices |
|  | epi/endo-zoic/phytic | Taxa that are attached to other living organisms |
|  | Attached | Taxa that are attached to hard substrates |
| Reproductive mode | Asexual | Taxa that reproduce asexually |
|  | Broadcast | Taxa that reproduce by releasing eggs and sperm in the water column |
|  | Brooder | Taxa that reproduce by internal or external fertilization but keep the eggs or larvae within the body, or in the burrow (parental care) or a body cavity until hatching |
|  | Benthic_Deposition | Taxa that produce egg sacs that are attached to a substrate |
| Larval development location | Planktotrophic | Larvae grow up planktonic and must feed on plankton for their development into their juvenile stage |
|  | Lecithotrophic | Planktonic development but initial food source is provided by the yolk from its egg |
|  | Benthic/Direct | No planktonic stage. Taxa develop fully within the sediment or brooded to juvenile |
| Skeleton | Soft | No exoskeleton, soft bodied |
|  | Calcified | Calcified exoskeleton |
|  | Chitin | Chitinous exoskeleton |
| Offspring size (μm) | ≤ 100 | Offspring smaller than 100 µm |
|  | >100 and ≤ 500 | Offspring between 100-500 µm |
|  | >500 and ≤ 1500 | Offspring between 500-1500 µm |
|  | >1500 | Offspring larger than 1500 µm |
| Offspring type | Juvenile | Offspring is released as juvenile |
|  | Larva | Offspring is released as larva |
|  | Egg | Offspring is released as egg |
| François, F., Poggiale, J.-C., Durbec, J.-P., & Stora, G. (1997). A New Approach for the Modelling of Sediment Reworking Induced by a Macrobenthic Community. *Acta Biotheoretica*, *45*(3), 295–319.<https://doi.org/10.1023/A:1000636109604>  Gardner, L. R., Sharma, P., & Moore, W. S. (1987). A regeneration model for the effect of bioturbation by fiddler crabs on 210Pb profiles in salt marsh sediments. *Journal of Environmental Radioactivity*, *5*(1), 25–36.<https://doi.org/10.1016/0265-931X(87)90042-7>  Solan, M., Cardinale, B. J., Downing, A. L., Engelhardt, K. A. M., Ruesink, J. L., & Srivastava, D. S. (2004). Extinction and  Ecosystem Function in the Marine Benthos. *Science*, *306*(5699), 1177–1180.<https://doi.org/10.1126/science.1103960> | | |

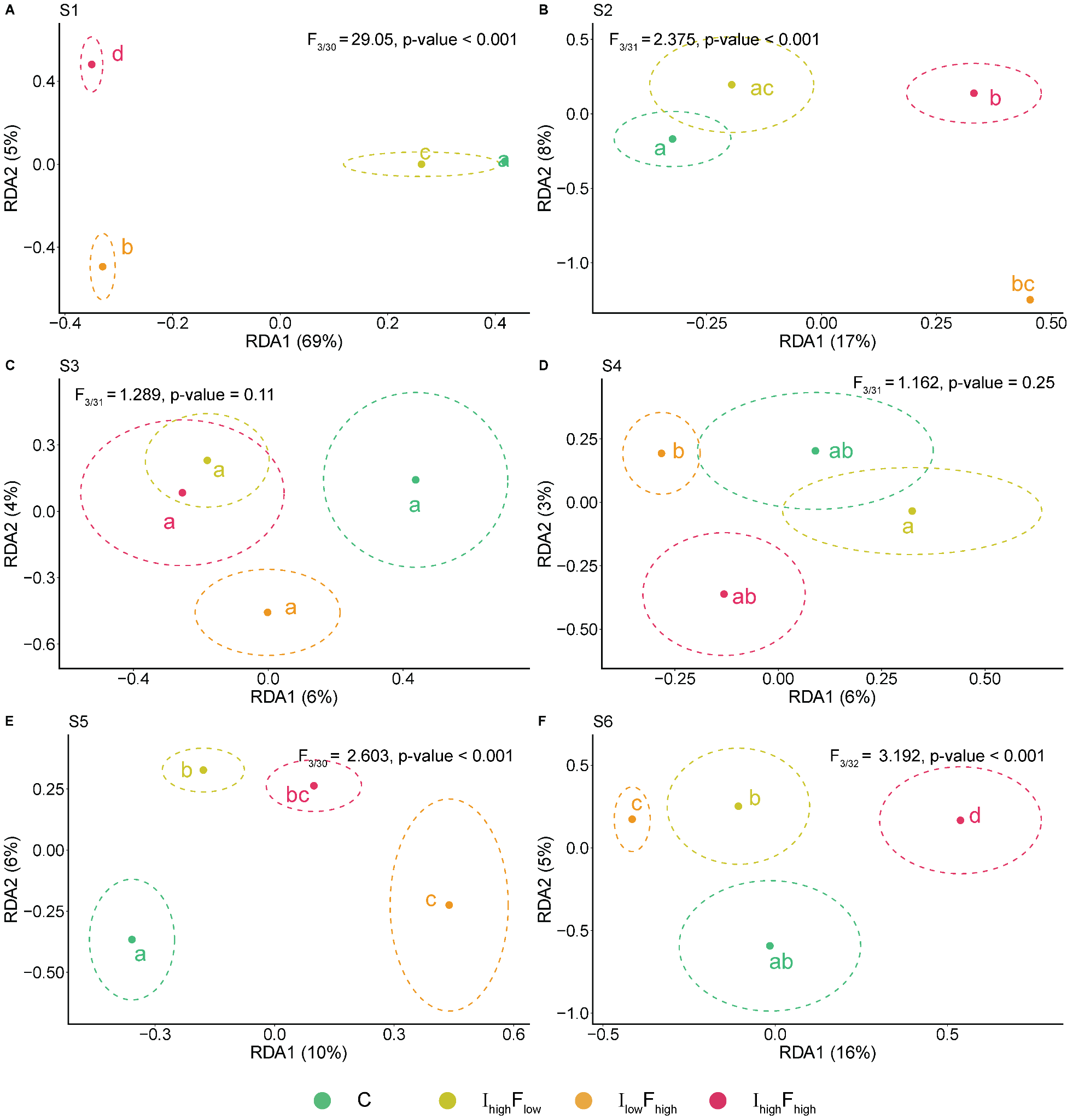
**Supplementary Figure S2.** Redundancy analysis of species composition at T0 (pre-disturbance). Axes represent the first two RDA axes. Each plot is a different site (see Figure 1B). Centroids for each treatment per time period are displayed with ellipses indicating the 95% confidence interval of the centroid. Significant differences between treatments are displayed through compact letter display. C is Control, I_high_F_low_ is High intensity, Low frequency, I_low_F_high_ is Low intensity, High frequency, and I_high_F_high_ is High intensity, High frequency.

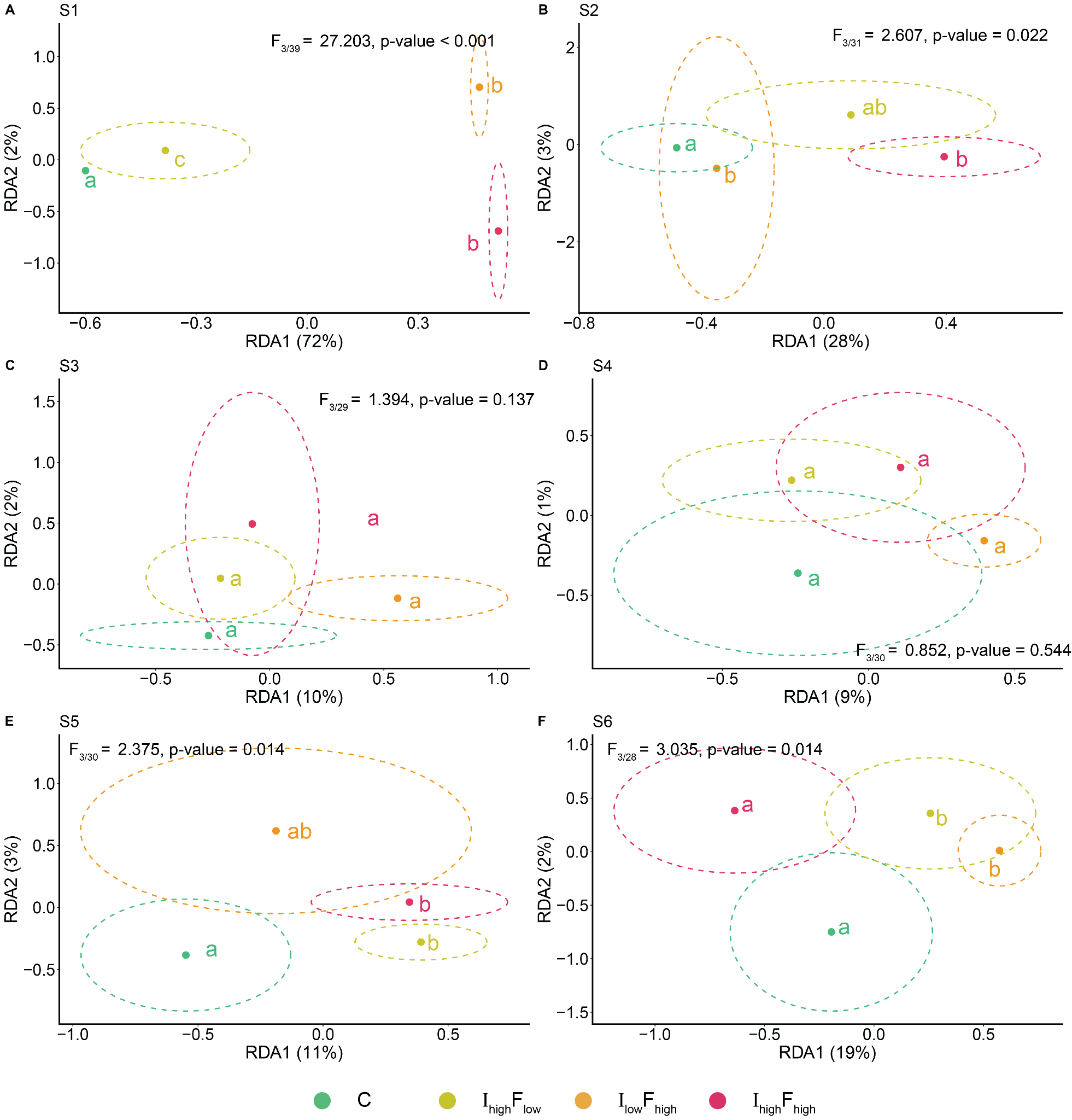
**Supplementary Figure S3.** Redundancy analysis of trait composition at T0 (pre-disturbance). Axes represent the first two RDA axes. Each plot is a different site (see Figure 1B). Centroids for each treatment per time period are displayed with ellipses indicating the 95% confidence interval of the centroid. Significant differences between treatments are displayed through compact letter display. C is Control, I_high_F_low_ is High intensity, Low frequency, I_low_F_high_ is Low intensity, High frequency, and I_high_F_high_ is High intensity, High frequency.

##### **Supplementary Table S3.** Bootstrapped coefficients for contrasts from all linear mixed models.

| **Response** | **Contrasts** | **Median** | **CI low** | **CI high** | **pd** | **p-value** |
| --- | --- | --- | --- | --- | --- | --- |
| **Evenness** | | | | | | |
|  | C - I_low_F_high_, T1 | 0.05 | -0.01 | 0.11 | 0.96 | 0.0780 |
|  | C - I_high_F_low_, T1 | -0.04 | -0.10 | 0.02 | 0.92 | 0.1650 |
|  | C - I_high_F_high_, T1 | -0.06 | -0.12 | 0.00 | 0.98 | 0.0406 |
|  | I_low_F_high_ - I_high_F_low_, T1 | -0.10 | -0.15 | -0.04 | 1.00 | 0.0014 |
|  | I_low_F_high_ - I_high_F_high_, T1 | -0.11 | -0.17 | -0.05 | 1.00 | 0.000 |
|  | I_high_F_low_ - I_high_F_high_, T1 | -0.02 | -0.07 | 0.04 | 0.74 | 0.5230 |
|  | C - I_low_F_high_, T3 | 0.00 | -0.06 | 0.06 | 0.52 | 0.9664 |
|  | C - I_high_F_low_, T3 | -0.04 | -0.10 | 0.01 | 0.93 | 0.1432 |
|  | C - I_high_F_high_, T3 | -0.01 | -0.06 | 0.05 | 0.59 | 0.8134 |
|  | I_low_F_high_ - I_high_F_low_, T3 | -0.04 | -0.10 | 0.01 | 0.92 | 0.1516 |
|  | I_low_F_high_ - I_high_F_high_, T3 | -0.01 | -0.07 | 0.05 | 0.61 | 0.7786 |
|  | I_high_F_low_ - I_high_F_high_, T3 | 0.03 | -0.02 | 0.09 | 0.89 | 0.2256 |
|  | C - I_low_F_high_, T6 | 0.10 | 0.04 | 0.16 | 1.00 | 0.006 |
|  | C - I_high_F_low_, T6 | 0.01 | -0.05 | 0.07 | 0.64 | 0.7164 |
|  | C - I_high_F_high_, T6 | 0.04 | -0.01 | 0.10 | 0.94 | 0.1218 |
|  | I_low_F_high_ - I_high_F_low_, T6 | -0.09 | -0.15 | -0.03 | 1.00 | 0.0022 |
|  | I_low_F_high_ - I_high_F_high_, T6 | -0.06 | -0.12 | 0.00 | 0.97 | 0.0552 |
|  | I_high_F_low_ - I_high_F_high_, T6 | 0.03 | -0.02 | 0.09 | 0.88 | 0.2386 |
|  | C - I_low_F_high_, T7 | 0.05 | -0.01 | 0.11 | 0.95 | 0.1052 |
|  | C - I_high_F_low_, T7 | -0.02 | -0.07 | 0.04 | 0.73 | 0.5476 |
|  | C - I_high_F_high_, T7 | -0.01 | -0.07 | 0.04 | 0.65 | 0.7066 |
|  | I_low_F_high_ - I_high_F_low_, T7 | -0.07 | -0.12 | -0.01 | 0.99 | 0.0282 |
|  | I_low_F_high_ - I_high_F_high_, T7 | -0.06 | -0.12 | 0.00 | 0.98 | 0.0438 |
|  | I_high_F_low_- I_high_F_high_, T7 | 0.01 | -0.05 | 0.06 | 0.59 | 0.8180 |
| **ENS2** | | | | | | |
|  | C - I_low_F_high_, T1 | 0.19 | 0.02 | 0.37 | 0.99 | 0.0278 |
|  | C - I_high_F_low_, T1 | -0.12 | -0.29 | 0.05 | 0.92 | 0.1610 |
|  | C - I_high_F_high_, T1 | -0.12 | -0.29 | 0.05 | 0.91 | 0.1826 |
|  | I_low_F_high_ - I_high_F_low_, T1 | -0.31 | -0.49 | -0.14 | 1.00 | 0.0002 |
|  | I_low_F_high_ - I_high_F_high_, T1 | -0.31 | -0.48 | -0.13 | 1.00 | 0.0002 |
|  | I_high_F_low_ - I_high_F_high_, T1 | 0.01 | -0.17 | 0.18 | 0.53 | 0.9432 |
|  | C - I_low_F_high_, T3 | 0.09 | -0.09 | 0.26 | 0.84 | 0.3232 |
|  | C - I_high_F_low_, T3 | -0.08 | -0.25 | 0.09 | 0.82 | 0.3696 |
|  | C - I_high_F_high_, T3 | 0.03 | -0.13 | 0.20 | 0.63 | 0.7328 |
|  | I_low_F_high_ - I_high_F_low_, T3 | -0.17 | -0.34 | 0.00 | 0.97 | 0.0544 |
|  | I_low_F_high_ - I_high_F_high_, T3 | -0.06 | -0.23 | 0.12 | 0.74 | 0.5182 |
|  | I_high_F_low_ - I_high_F_high_, T3 | 0.11 | -0.06 | 0.28 | 0.90 | 0.2020 |
|  | C - I_low_F_high_, T6 | 0.30 | 0.13 | 0.47 | 1.00 | 0.0006 |
|  | C - I_high_F_low_, T6 | 0.05 | -0.12 | 0.23 | 0.72 | 0.5524 |
|  | C - I_high_F_high_, T6 | 0.11 | -0.06 | 0.28 | 0.90 | 0.1902 |
|  | I_low_F_high_ - I_high_F_low_, T6 | -0.25 | -0.42 | -0.07 | 1.00 | 0.0054 |
|  | I_low_F_high_ - I_high_F_high_, T6 | -0.19 | -0.36 | -0.02 | 0.98 | 0.0340 |
|  | I_high_F_low_ - I_high_F_high_, T6 | 0.06 | -0.11 | 0.23 | 0.766 | 0.4814 |
|  | C - I_low_F_high_, T7 | 0.08 | -0.09 | 0.25 | 0.82 | 0.3650 |
|  | C - I_high_F_low_, T7 | -0.06 | -0.23 | 0.11 | 0.75 | 0.4938 |
|  | C - I_high_F_high_, T7 | -0.13 | -0.30 | 0.03 | 0.94 | 0.1220 |
|  | I_low_F_high_ - I_high_F_low_, T7 | -0.14 | -0.32 | 0.04 | 0.94 | 0.1194 |
|  | I_low_F_high_- I_high_F_high_, T7 | -0.21 | -0.39 | -0.04 | 0.99 | 0.0134 |
|  | I_high_F_low_- I_high_F_high_, T7 | -0.07 | -0.24 | 0.10 | 0.79 | 0.4176 |
| **Long-living species** | | | | | | |
|  | C - I_low_F_high_, T1 | 0.00 | -0.06 | 0.06 | 0.52 | 0.9598 |
|  | C - I_high_F_low_, T1 | -0.01 | -0.06 | 0.05 | 0.58 | 0.8330 |
|  | C - I_high_F_high_, T1 | 0.01 | -0.04 | 0.07 | 0.66 | 0.6804 |
|  | I_low_F_high_ - I_high_F_low_, T1 | -0.01 | -0.06 | 0.05 | 0.59 | 0.8276 |
|  | I_low_F_high_ - I_high_F_high_, T1 | 0.01 | -0.05 | 0.07 | 0.64 | 0.7156 |
|  | I_high_F_low_ - I_high_F_high_, T1 | 0.02 | -0.04 | 0.07 | 0.72 | 0.5524 |
|  | C - I_low_F_high_, T3 | 0.02 | -0.04 | 0.08 | 0.76 | 0.4704 |
|  | C - I_high_F_low_, T3 | 0.01 | -0.04 | 0.07 | 0.69 | 0.6194 |
|  | C - I_high_F_high_, T3 | 0.1 | 0.04 | 0.16 | 1 | 0.0002 |
|  | I_low_F_high_ - I_high_F_low_, T3 | -0.01 | -0.06 | 0.05 | 0.6 | 0.8042 |
|  | I_low_F_high_ - I_high_F_high_, T3 | 0.08 | 0.02 | 0.14 | 1 | 0.005 |
|  | I_high_F_low_ - I_high_F_high_, T3 | 0.09 | 0.03 | 0.14 | 1 | 0.0028 |
|  | C - I_low_F_high_, T6 | 0.08 | 0.02 | 0.14 | 1 | 0.0062 |
|  | C - I_high_F_low_, T6 | 0.02 | -0.03 | 0.08 | 0.8 | 0.3908 |
|  | C - I_high_F_high_, T6 | 0.1 | 0.04 | 0.15 | 1 | 0.0002 |
|  | I_low_F_high_ - I_high_F_low_, T6 | -0.06 | -0.11 | 0 | 0.98 | 0.0488 |
|  | I_low_F_high_ - I_high_F_high_, T6 | 0.02 | -0.04 | 0.07 | 0.72 | 0.551 |
|  | I_high_F_low_ - I_high_F_high_, T6 | 0.07 | 0.02 | 0.13 | 1 | 0.0098 |
|  | C - I_low_F_high_, T7 | 0.07 | 0.01 | 0.13 | 0.99 | 0.0158 |
|  | C - I_high_F_low_, T7 | 0.01 | -0.04 | 0.07 | 0.66 | 0.6702 |
|  | C - I_high_F_high_, T7 | 0.06 | 0.01 | 0.12 | 0.99 | 0.0228 |
|  | I_low_F_high_ - I_high_F_low_, T7 | -0.06 | -0.12 | 0 | 0.98 | 0.047 |
|  | I_low_F_high_ - I_high_F_high_, T7 | -0.01 | -0.06 | 0.05 | 0.6 | 0.7906 |
|  | I_high_F_low_- I_high_F_high_, T7 | 0.05 | -0.01 | 0.11 | 0.96 | 0.0756 |
| **Rao's Q** | | | | | | |
|  | C - I_low_F_high_, T1 | 0.52 | 0.21 | 0.82 | 1 | 0.0014 |
|  | C - I_high_F_low_, T1 | 0.06 | -0.24 | 0.36 | 0.64 | 0.7128 |
|  | C - I_high_F_high_, T1 | 0.23 | -0.06 | 0.53 | 0.94 | 0.1228 |
|  | I_low_F_high_ - I_high_F_low_, T1 | -0.46 | -0.76 | -0.14 | 1 | 0.0044 |
|  | I_low_F_high_ - I_high_F_high_, T1 | -0.28 | -0.59 | 0.02 | 0.97 | 0.0692 |
|  | I_high_F_low_ - I_high_F_high_, T1 | 0.18 | -0.12 | 0.47 | 0.88 | 0.2398 |
|  | C - I_low_F_high_, T3 | 0.42 | 0.11 | 0.72 | 1 | 0.0068 |
|  | C - I_high_F_low_, T3 | 0.22 | -0.07 | 0.51 | 0.93 | 0.1418 |
|  | C - I_high_F_high_, T3 | 0.36 | 0.07 | 0.65 | 0.99 | 0.0160 |
|  | I_low_F_high_ - I_high_F_low_, T3 | -0.20 | -0.49 | 0.10 | 0.90 | 0.1948 |
|  | I_low_F_high_ - I_high_F_high_, T3 | -0.06 | -0.36 | 0.24 | 0.65 | 0.6912 |
|  | I_high_F_low_ - I_high_F_high_, T3 | 0.14 | -0.16 | 0.43 | 0.82 | 0.3638 |
|  | C - I_low_F_high_, T6 | 0.50 | 0.19 | 0.80 | 1 | 0.0014 |
|  | C - I_high_F_low_, T6 | 0.07 | -0.22 | 0.36 | 0.68 | 0.6442 |
|  | C - I_high_F_high_, T6 | 0.33 | 0.05 | 0.62 | 0.99 | 0.0202 |
|  | I_low_F_high_ - I_high_F_low_, T6 | -0.43 | -0.73 | -0.13 | 1 | 0.0048 |
|  | I_low_F_high_ - I_high_F_high_, T6 | -0.16 | -0.46 | 0.14 | 0.86 | 0.2712 |
|  | I_high_F_low_ - I_high_F_high_, T6 | 0.27 | -0.03 | 0.56 | 0.96 | 0.0722 |
|  | C - I_low_F_high_, T7 | 0.36 | 0.05 | 0.66 | 0.99 | 0.0212 |
|  | C - I_high_F_low_, T7 | -0.04 | -0.34 | 0.25 | 0.61 | 0.7896 |
|  | C - I_high_F_high_, T7 | 0.29 | 0.00 | 0.58 | 0.98 | 0.0462 |
|  | I_low_F_high_ - I_high_F_low_, T7 | -0.40 | -0.71 | -0.09 | 0.99 | 0.0116 |
|  | I_low_F_high_ - I_high_F_high_, T7 | -0.06 | -0.36 | 0.23 | 0.66 | 0.6724 |
|  | I_high_F_low_- I_high_F_high_, T7 | 0.33 | 0.04 | 0.63 | 0.99 | 0.0268 |
| **Scavengers and predators** | | | | | | |
|  | C - I_low_F_high_, T1 | -0.12 | -0.17 | -0.06 | 1 | 0 |
|  | C - I_high_F_low_, T1 | -0.06 | -0.12 | -0.01 | 0.99 | 0.0216 |
|  | C - I_high_F_high_, T1 | -0.1 | -0.16 | -0.05 | 1 | 0 |
|  | I_low_F_high_ - I_high_F_low_, T1 | 0.05 | 0 | 0.11 | 0.97 | 0.056 |
|  | I_low_F_high_ - I_high_F_high_, T1 | 0.01 | -0.04 | 0.07 | 0.67 | 0.6526 |
|  | I_high_F_low_ - I_high_F_high_, T1 | -0.04 | -0.09 | 0.01 | 0.94 | 0.1242 |
|  | C - I_low_F_high_, T3 | -0.12 | -0.17 | -0.06 | 1 | 0 |
|  | C - I_high_F_low_, T3 | -0.03 | -0.08 | 0.02 | 0.88 | 0.2454 |
|  | C - I_high_F_high_, T3 | -0.09 | -0.15 | -0.04 | 1 | 0.0006 |
|  | I_low_F_high_ - I_high_F_low_, T3 | 0.08 | 0.03 | 0.14 | 1 | 0.0016 |
|  | I_low_F_high_ - I_high_F_high_, T3 | 0.02 | -0.03 | 0.08 | 0.8 | 0.401 |
|  | I_high_F_low_ - I_high_F_high_, T3 | -0.06 | -0.11 | -0.01 | 0.99 | 0.0202 |
|  | C - I_low_F_high_, T6 | -0.06 | -0.11 | 0 | 0.98 | 0.0348 |
|  | C - I_high_F_low_, T6 | -0.01 | -0.06 | 0.05 | 0.58 | 0.8364 |
|  | C - I_high_F_high_, T6 | -0.07 | -0.12 | -0.02 | 1 | 0.0074 |
|  | I_low_F_high_ - I_high_F_low_, T6 | 0.05 | 0 | 0.11 | 0.97 | 0.0608 |
|  | I_low_F_high_ - I_high_F_high_, T6 | -0.01 | -0.07 | 0.04 | 0.68 | 0.6354 |
|  | I_high_F_low_ - I_high_F_high_, T6 | -0.06 | -0.12 | -0.01 | 0.99 | 0.017 |
|  | C - I_low_F_high_, T7 | -0.05 | -0.1 | 0.01 | 0.96 | 0.0736 |
|  | C - I_high_F_low_, T7 | 0 | -0.05 | 0.05 | 0.51 | 0.9888 |
|  | C - I_high_F_high_, T7 | -0.07 | -0.13 | -0.02 | 1 | 0.0054 |
|  | I_low_F_high_ - I_high_F_low_, T7 | 0.05 | 0 | 0.1 | 0.96 | 0.0706 |
|  | I_low_F_high_ - I_high_F_high_, T7 | -0.02 | -0.08 | 0.03 | 0.8 | 0.398 |
|  | I_high_F_low_- I_high_F_high_, T7 | -0.07 | -0.13 | -0.02 | 1 | 0.0064 |
| **ENS1** | | | | | | |
|  | C - I_low_F_high_, T1 | 0.25 | 0.06 | 0.43 | 1.00 | 0.0086 |
|  | C - I_high_F_low_, T1 | -0.07 | -0.25 | 0.10 | 0.79 | 0.4278 |
|  | C - I_high_F_high_, T1 | -0.04 | -0.22 | 0.14 | 0.69 | 0.6174 |
|  | I_low_F_high_ - I_high_F_low_, T1 | -0.32 | -0.50 | -0.14 | 1.00 | 0.0004 |
|  | I_low_F_high_ - I_high_F_high_, T1 | -0.29 | -0.48 | -0.10 | 1.00 | 0.0024 |
|  | I_high_F_low_ - I_high_F_high_, T1 | 0.03 | -0.15 | 0.21 | 0.62 | 0.7630 |
|  | C - I_low_F_high_, T3 | 0.15 | -0.03 | 0.33 | 0.95 | 0.1008 |
|  | C - I_high_F_low_, T3 | -0.02 | -0.19 | 0.16 | 0.58 | 0.8358 |
|  | C - I_high_F_high_, T3 | 0.10 | -0.07 | 0.28 | 0.88 | 0.2464 |
|  | I_low_F_high_ - I_high_F_low_, T3 | -0.17 | -0.35 | 0.01 | 0.97 | 0.0646 |
|  | I_low_F_high_ - I_high_F_high_, T3 | -0.05 | -0.23 | 0.14 | 0.70 | 0.5946 |
|  | I_high_F_low_ - I_high_F_high_, T3 | 0.12 | -0.05 | 0.29 | 0.92 | 0.1686 |
|  | C - I_low_F_high_, T6 | 0.30 | 0.12 | 0.48 | 1.00 | 0.0016 |
|  | C - I_high_F_low_, T6 | 0.07 | -0.11 | 0.24 | 0.77 | 0.4668 |
|  | C - I_high_F_high_, T6 | 0.14 | -0.04 | 0.31 | 0.94 | 0.1222 |
|  | I_low_F_high_ - I_high_F_low_, T6 | -0.24 | -0.42 | -0.06 | 1.00 | 0.0098 |
|  | I_low_F_high_ - I_high_F_high_, T6 | -0.17 | -0.34 | 0.01 | 0.97 | 0.0672 |
|  | I_high_F_low_ - I_high_F_high_, T6 | 0.07 | -0.10 | 0.25 | 0.79 | 0.4192 |
|  | C - I_low_F_high_, T7 | 0.13 | -0.06 | 0.31 | 0.92 | 0.1688 |
|  | C - I_high_F_low_, T7 | 0.00 | -0.19 | 0.18 | 0.50 | 0.9990 |
|  | C - I_high_F_high_, T7 | -0.04 | -0.22 | 0.13 | 0.68 | 0.6380 |
|  | I_low_F_high_ - I_high_F_low_, T7 | -0.13 | -0.31 | 0.05 | 0.92 | 0.1628 |
|  | I_low_F_high_ - I_high_F_high_, T7 | -0.17 | -0.35 | 0.01 | 0.97 | 0.0646 |
|  | I_high_F_low_- I_high_F_high_, T7 | -0.04 | -0.22 | 0.14 | 0.67 | 0.6532 |
| **Species richness** | | | | | | |
|  | C - I_low_F_high_, T1 | 0.09 | -0.09 | 0.28 | 0.85 | 0.3026 |
|  | C - I_high_F_low_, T1 | 0.05 | -0.13 | 0.22 | 0.7 | 0.6028 |
|  | C - I_high_F_high_, T1 | 0.12 | -0.06 | 0.3 | 0.9 | 0.1938 |
|  | I_low_F_high_ - I_high_F_low_, T1 | -0.05 | -0.23 | 0.13 | 0.7 | 0.5966 |
|  | I_low_F_high_ - I_high_F_high_, T1 | 0.02 | -0.16 | 0.21 | 0.59 | 0.8288 |
|  | I_high_F_low_ - I_high_F_high_, T1 | 0.07 | -0.11 | 0.25 | 0.77 | 0.4518 |
|  | C - I_low_F_high_, T3 | 0.15 | -0.03 | 0.33 | 0.95 | 0.1036 |
|  | C - I_high_F_low_, T3 | 0.13 | -0.04 | 0.31 | 0.93 | 0.138 |
|  | C - I_high_F_high_, T3 | 0.14 | -0.03 | 0.32 | 0.95 | 0.1064 |
|  | I_low_F_high_ - I_high_F_low_, T3 | -0.02 | -0.2 | 0.17 | 0.57 | 0.8522 |
|  | I_low_F_high_ - I_high_F_high_, T3 | -0.01 | -0.19 | 0.17 | 0.54 | 0.923 |
|  | I_high_F_low_ - I_high_F_high_, T3 | 0.01 | -0.17 | 0.18 | 0.53 | 0.9388 |
|  | C - I_low_F_high_, T6 | 0.01 | -0.17 | 0.2 | 0.55 | 0.8954 |
|  | C - I_high_F_low_, T6 | 0.04 | -0.14 | 0.21 | 0.68 | 0.6396 |
|  | C - I_high_F_high_, T6 | -0.01 | -0.18 | 0.17 | 0.52 | 0.9522 |
|  | I_low_F_high_ - I_high_F_low_, T6 | 0.03 | -0.15 | 0.21 | 0.62 | 0.7616 |
|  | I_low_F_high_ - I_high_F_high_, T6 | -0.02 | -0.2 | 0.16 | 0.58 | 0.8432 |
|  | I_high_F_low_ - I_high_F_high_, T6 | -0.05 | -0.22 | 0.13 | 0.7 | 0.5982 |
|  | C - I_low_F_high_, T7 | -0.02 | -0.2 | 0.16 | 0.6 | 0.8054 |
|  | C - I_high_F_low_, T7 | 0.06 | -0.12 | 0.23 | 0.74 | 0.5292 |
|  | C - I_high_F_high_, T7 | -0.07 | -0.24 | 0.11 | 0.77 | 0.4548 |
|  | I_low_F_high_ - I_high_F_low_, T7 | 0.08 | -0.1 | 0.26 | 0.8 | 0.4038 |
|  | I_low_F_high_ - I_high_F_high_, T7 | -0.05 | -0.22 | 0.14 | 0.68 | 0.6378 |
|  | I_high_F_low_- I_high_F_high_, T7 | -0.12 | -0.3 | 0.06 | 0.91 | 0.1822 |
| **Suspension feeder** | | | | | | |
|  | C - I_low_F_high_, T1 | 0.03 | -0.04 | 0.09 | 0.78 | 0.4432 |
|  | C - I_high_F_low_, T1 | -0.01 | -0.08 | 0.05 | 0.65 | 0.7022 |
|  | C - I_high_F_high_, T1 | 0 | -0.07 | 0.07 | 0.52 | 0.965 |
|  | I_low_F_high_ - I_high_F_low_, T1 | -0.04 | -0.11 | 0.03 | 0.87 | 0.2546 |
|  | I_low_F_high_ - I_high_F_high_, T1 | -0.03 | -0.1 | 0.04 | 0.79 | 0.4226 |
|  | I_high_F_low_ - I_high_F_high_, T1 | 0.01 | -0.06 | 0.08 | 0.63 | 0.7474 |
|  | C - I_low_F_high_, T3 | 0.08 | 0.01 | 0.15 | 0.99 | 0.0198 |
|  | C - I_high_F_low_, T3 | 0.05 | -0.01 | 0.12 | 0.95 | 0.1014 |
|  | C - I_high_F_high_, T3 | 0.13 | 0.07 | 0.2 | 1 | 0.0002 |
|  | I_low_F_high_ - I_high_F_low_, T3 | -0.03 | -0.1 | 0.04 | 0.79 | 0.4292 |
|  | I_low_F_high_ - I_high_F_high_, T3 | 0.05 | -0.01 | 0.12 | 0.94 | 0.1232 |
|  | I_high_F_low_ - I_high_F_high_, T3 | 0.08 | 0.02 | 0.14 | 0.99 | 0.0166 |
|  | C - I_low_F_high_, T6 | 0.07 | 0 | 0.13 | 0.97 | 0.0578 |
|  | C - I_high_F_low_, T6 | -0.03 | -0.09 | 0.04 | 0.79 | 0.4274 |
|  | C - I_high_F_high_, T6 | 0.07 | 0.01 | 0.14 | 0.99 | 0.0258 |
|  | I_low_F_high_ - I_high_F_low_, T6 | -0.09 | -0.16 | -0.03 | 1 | 0.007 |
|  | I_low_F_high_ - I_high_F_high_, T6 | 0.01 | -0.06 | 0.07 | 0.59 | 0.8234 |
|  | I_high_F_low_ - I_high_F_high_, T6 | 0.1 | 0.04 | 0.17 | 1 | 0.0026 |
|  | C - I_low_F_high_, T7 | 0.08 | 0.01 | 0.14 | 0.99 | 0.0274 |
|  | C - I_high_F_low_, T7 | 0.01 | -0.05 | 0.08 | 0.66 | 0.6714 |
|  | C - I_high_F_high_, T7 | 0.05 | -0.01 | 0.12 | 0.94 | 0.1138 |
|  | I_low_F_high_ - I_high_F_low_, T7 | -0.06 | -0.13 | 0 | 0.97 | 0.065 |
|  | I_low_F_high_ - I_high_F_high_, T7 | -0.02 | -0.09 | 0.04 | 0.77 | 0.4658 |
|  | I_high_F_low_ - I_high_F_high_, T7 | 0.04 | -0.03 | 0.1 | 0.87 | 0.254 |
| **Hellinger distance taxonomic community within treatment** | | | | | | |
|  | C - I_low_F_high_, T0 | -0.03 | -0.1 | 0.04 | 0.82 | 0.3676 |
|  | C - I_high_F_low_, T0 | -0.03 | -0.1 | 0.03 | 0.85 | 0.2954 |
|  | C - I_high_F_high_, T0 | -0.06 | -0.12 | 0.01 | 0.96 | 0.0796 |
|  | I_low_F_high_ - I_high_F_low_, T0 | 0 | -0.07 | 0.06 | 0.54 | 0.916 |
|  | I_low_F_high_ - I_high_F_high_, T0 | -0.03 | -0.09 | 0.04 | 0.78 | 0.4316 |
|  | I_high_F_low_ - I_high_F_high_, T0 | -0.02 | -0.09 | 0.04 | 0.75 | 0.493 |
|  | C - I_low_F_high_, T1 | 0.03 | -0.03 | 0.1 | 0.83 | 0.3406 |
|  | C - I_high_F_low_, T1 | -0.04 | -0.11 | 0.02 | 0.9 | 0.2018 |
|  | C - I_high_F_high_, T1 | -0.18 | -0.25 | -0.12 | 1 | 0 |
|  | I_low_F_high_ - I_high_F_low_, T1 | -0.07 | -0.14 | -0.01 | 0.99 | 0.0282 |
|  | I_low_F_high_ - I_high_F_high_, T1 | -0.21 | -0.28 | -0.15 | 1 | 0 |
|  | I_high_F_low_ - I_high_F_high_, T1 | -0.14 | -0.21 | -0.08 | 1 | 0 |
|  | C - I_low_F_high_, T3 | 0.02 | -0.05 | 0.08 | 0.73 | 0.546 |
|  | C - I_high_F_low_, T3 | -0.03 | -0.09 | 0.04 | 0.79 | 0.4134 |
|  | C - I_high_F_high_, T3 | -0.01 | -0.07 | 0.06 | 0.59 | 0.8154 |
|  | I_low_F_high_ - I_high_F_low_, T3 | -0.05 | -0.11 | 0.02 | 0.92 | 0.1542 |
|  | I_low_F_high_ - I_high_F_high_, T3 | -0.03 | -0.09 | 0.04 | 0.79 | 0.421 |
|  | I_high_F_low_ - I_high_F_high_, T3 | 0.02 | -0.04 | 0.08 | 0.73 | 0.5346 |
|  | C - I_low_F_high_, T6 | 0.03 | -0.04 | 0.09 | 0.78 | 0.4312 |
|  | C - I_high_F_low_, T6 | -0.03 | -0.09 | 0.04 | 0.78 | 0.4396 |
|  | C - I_high_F_high_, T6 | -0.07 | -0.14 | -0.01 | 0.99 | 0.0274 |
|  | I_low_F_high_ - I_high_F_low_, T6 | -0.05 | -0.11 | 0.01 | 0.94 | 0.1116 |
|  | I_low_F_high_ - I_high_F_high_, T6 | -0.1 | -0.16 | -0.04 | 1 | 0.0028 |
|  | I_high_F_low_ - I_high_F_high_, T6 | -0.05 | -0.11 | 0.02 | 0.93 | 0.1358 |
|  | C - I_low_F_high_, T7 | 0.03 | -0.03 | 0.1 | 0.86 | 0.2798 |
|  | C - I_high_F_low_, T7 | -0.11 | -0.17 | -0.05 | 1 | 0.001 |
|  | C - I_high_F_high_, T7 | -0.1 | -0.17 | -0.04 | 1 | 0.0016 |
|  | I_low_F_high_ - I_high_F_low_, T7 | -0.14 | -0.21 | -0.08 | 1 | 0 |
|  | I_low_F_high_ - I_high_F_high_, T7 | -0.14 | -0.2 | -0.08 | 1 | 0 |
|  | I_high_F_low_- I_high_F_high_, T7 | 0 | -0.06 | 0.07 | 0.54 | 0.9216 |
| **Euclidean distance trait community within treatment** | | | | | | |
|  | C - I_low_F_high_, T0 | -0.07 | -0.24 | 0.09 | 0.8 | 0.3928 |
|  | C - I_high_F_low_, T0 | -0.08 | -0.24 | 0.09 | 0.83 | 0.3362 |
|  | C - I_high_F_high_, T0 | -0.13 | -0.29 | 0.03 | 0.95 | 0.104 |
|  | I_low_F_high_ - I_high_F_low_, T0 | -0.01 | -0.17 | 0.16 | 0.53 | 0.9422 |
|  | I_low_F_high_ - I_high_F_high_, T0 | -0.06 | -0.22 | 0.1 | 0.76 | 0.4708 |
|  | I_high_F_low_ - I_high_F_high_, T0 | -0.06 | -0.21 | 0.11 | 0.75 | 0.5076 |
|  | C - I_low_F_high_, T1 | 0.02 | -0.14 | 0.18 | 0.6 | 0.8072 |
|  | C - I_high_F_low_, T1 | -0.01 | -0.18 | 0.15 | 0.56 | 0.8718 |
|  | C - I_high_F_high_, T1 | -0.47 | -0.63 | -0.31 | 1 | 0 |
|  | I_low_F_high_ - I_high_F_low_, T1 | -0.04 | -0.19 | 0.12 | 0.66 | 0.6738 |
|  | I_low_F_high_ - I_high_F_high_, T1 | -0.49 | -0.66 | -0.33 | 1 | 0 |
|  | I_high_F_low_ - I_high_F_high_, T1 | -0.46 | -0.62 | -0.29 | 1 | 0 |
|  | C - I_low_F_high_, T3 | 0.11 | -0.05 | 0.27 | 0.91 | 0.1712 |
|  | C - I_high_F_low_, T3 | -0.04 | -0.2 | 0.11 | 0.71 | 0.582 |
|  | C - I_high_F_high_, T3 | -0.1 | -0.25 | 0.07 | 0.88 | 0.2486 |
|  | I_low_F_high_ - I_high_F_low_, T3 | -0.15 | -0.31 | 0 | 0.97 | 0.0512 |
|  | I_low_F_high_ - I_high_F_high_, T3 | -0.21 | -0.36 | -0.05 | 0.99 | 0.0128 |
|  | I_high_F_low_ - I_high_F_high_, T3 | -0.05 | -0.21 | 0.11 | 0.73 | 0.5374 |
|  | C - I_low_F_high_, T6 | -0.1 | -0.26 | 0.06 | 0.89 | 0.212 |
|  | C - I_high_F_low_, T6 | -0.19 | -0.35 | -0.03 | 0.99 | 0.0188 |
|  | C - I_high_F_high_, T6 | -0.35 | -0.51 | -0.19 | 1 | 0 |
|  | I_low_F_high_ - I_high_F_low_, T6 | -0.09 | -0.24 | 0.07 | 0.87 | 0.2682 |
|  | I_low_F_high_ - I_high_F_high_, T6 | -0.24 | -0.4 | -0.09 | 1 | 0.0022 |
|  | I_high_F_low_ - I_high_F_high_, T6 | -0.16 | -0.31 | 0 | 0.97 | 0.0536 |
|  | C - I_low_F_high_, T7 | 0.13 | -0.02 | 0.29 | 0.95 | 0.1028 |
|  | C - I_high_F_low_, T7 | -0.18 | -0.34 | -0.02 | 0.99 | 0.0244 |
|  | C - I_high_F_high_, T7 | -0.14 | -0.3 | 0.02 | 0.95 | 0.0944 |
|  | I_low_F_high_ - I_high_F_low_, T7 | -0.32 | -0.48 | -0.16 | 1 | 0 |
|  | I_low_F_high_ - I_high_F_high_, T7 | -0.27 | -0.43 | -0.11 | 1 | 0.0012 |
|  | I_high_F_low_- I_high_F_high_, T7 | 0.05 | -0.11 | 0.21 | 0.72 | 0.5656 |
| **Hellinger distance taxonomic community turnover** | | | | | | |
|  | C - I_low_F_high_, T1 | -0.29 | -0.42 | -0.17 | 1 | 0 |
|  | C - I_high_F_low_, T1 | -0.09 | -0.21 | 0.03 | 0.92 | 0.1566 |
|  | C - I_high_F_high_, T1 | -0.29 | -0.41 | -0.17 | 1 | 0 |
|  | I_low_F_high_- I_high_F_low_, T1 | 0.21 | 0.08 | 0.33 | 1 | 0.0014 |
|  | I_low_F_high_ - I_high_F_high_, T1 | 0 | -0.12 | 0.13 | 0.52 | 0.955 |
|  | I_high_F_low_ - I_high_F_high_, T1 | -0.2 | -0.32 | -0.08 | 1 | 0.0008 |
|  | C - I_low_F_high_, T3 | -0.14 | -0.26 | -0.01 | 0.99 | 0.028 |
|  | C - I_high_F_low_, T3 | 0.04 | -0.08 | 0.16 | 0.73 | 0.5318 |
|  | C - I_high_F_high_, T3 | -0.13 | -0.24 | -0.01 | 0.98 | 0.0332 |
|  | I_low_F_high_ - I_high_F_low_, T3 | 0.17 | 0.05 | 0.3 | 1 | 0.0056 |
|  | I_low_F_high_ - I_high_F_high_, T3 | 0.01 | -0.11 | 0.13 | 0.56 | 0.888 |
|  | I_high_F_low_ - I_high_F_high_, T3 | -0.16 | -0.28 | -0.05 | 1 | 0.005 |
|  | C - I_low_F_high_, T6 | -0.11 | -0.23 | 0.01 | 0.97 | 0.0692 |
|  | C - I_high_F_low_, T6 | -0.04 | -0.16 | 0.08 | 0.75 | 0.491 |
|  | C - I_high_F_high_, T6 | -0.13 | -0.25 | -0.02 | 0.99 | 0.0254 |
|  | I_low_F_high_ - I_high_F_low_, T6 | 0.07 | -0.05 | 0.19 | 0.87 | 0.2534 |
|  | I_low_F_high_ - I_high_F_high_, T6 | -0.02 | -0.14 | 0.1 | 0.61 | 0.7726 |
|  | I_high_F_low_ - I_high_F_high_, T6 | -0.09 | -0.21 | 0.03 | 0.93 | 0.1444 |
|  | C - I_low_F_high_, T7 | -0.1 | -0.23 | 0.02 | 0.95 | 0.0924 |
|  | C - I_high_F_low_, T7 | -0.1 | -0.22 | 0.02 | 0.96 | 0.0844 |
|  | C - I_high_F_high_, T7 | -0.22 | -0.34 | -0.1 | 1 | 0 |
|  | I_low_F_high_ - I_high_F_low_, T7 | 0 | -0.12 | 0.12 | 0.5 | 0.9952 |
|  | I_low_F_high_ - I_high_F_high_, T7 | -0.12 | -0.24 | 0.01 | 0.97 | 0.0634 |
|  | I_high_F_low_- I_high_F_high_, T7 | -0.12 | -0.24 | 0 | 0.97 | 0.0536 |
| **Euclidean distance trait community turnover** | | | | | | |
|  | C - I_low_F_high_, T1 | -0.72 | -1.04 | -0.4 | 1 | 0 |
|  | C - I_high_F_low_, T1 | -0.25 | -0.55 | 0.07 | 0.95 | 0.1098 |
|  | C - I_high_F_high_, T1 | -0.73 | -1.04 | -0.43 | 1 | 0 |
|  | I_low_F_high_ - I_high_F_low_, T1 | 0.47 | 0.16 | 0.79 | 1 | 0.0032 |
|  | I_low_F_high_ - I_high_F_high_, T1 | -0.01 | -0.32 | 0.31 | 0.52 | 0.9636 |
|  | I_high_F_low_ - I_high_F_high_, T1 | -0.48 | -0.79 | -0.17 | 1 | 0.0028 |
|  | C - I_low_F_high_, T3 | -0.15 | -0.47 | 0.16 | 0.83 | 0.3366 |
|  | C - I_high_F_low_, T3 | 0.16 | -0.14 | 0.46 | 0.84 | 0.321 |
|  | C - I_high_F_high_, T3 | -0.21 | -0.51 | 0.09 | 0.91 | 0.1756 |
|  | I_low_F_high_ - I_high_F_low_, T3 | 0.31 | -0.01 | 0.62 | 0.97 | 0.056 |
|  | I_low_F_high_ - I_high_F_high_, T3 | -0.06 | -0.37 | 0.25 | 0.64 | 0.7136 |
|  | I_high_F_low_ - I_high_F_high_, T3 | -0.36 | -0.67 | -0.07 | 0.99 | 0.0174 |
|  | C - I_low_F_high_, T6 | -0.35 | -0.66 | -0.04 | 0.99 | 0.0266 |
|  | C - I_high_F_low_, T6 | -0.13 | -0.44 | 0.16 | 0.8 | 0.392 |
|  | C - I_high_F_high_, T6 | -0.38 | -0.67 | -0.08 | 0.99 | 0.0168 |
|  | I_low_F_high_ - I_high_F_low_, T6 | 0.22 | -0.09 | 0.52 | 0.91 | 0.1744 |
|  | I_low_F_high_ - I_high_F_high_, T6 | -0.03 | -0.34 | 0.28 | 0.57 | 0.8656 |
|  | I_high_F_low_ - I_high_F_high_, T6 | -0.24 | -0.53 | 0.06 | 0.94 | 0.112 |
|  | C - I_low_F_high_, T7 | -0.31 | -0.61 | 0 | 0.97 | 0.0536 |
|  | C - I_high_F_low_, T7 | -0.31 | -0.62 | -0.01 | 0.98 | 0.0428 |
|  | C - I_high_F_high_, T7 | -0.59 | -0.88 | -0.28 | 1 | 0 |
|  | I_low_F_high_ - I_high_F_low_, T7 | 0 | -0.31 | 0.31 | 0.51 | 0.9806 |
|  | I_low_F_high_ - I_high_F_high_, T7 | -0.28 | -0.59 | 0.03 | 0.96 | 0.0792 |
|  | I_high_F_low_ - I_high_F_high_, T7 | -0.27 | -0.57 | 0.03 | 0.96 | 0.0764 |
